# Genome-wide association study for age-related hearing loss in CFW mice

**DOI:** 10.1101/2024.06.10.598304

**Authors:** Oksana Polesskaya, Ely Boussaty, Riyan Cheng, Olivia Lamonte, Thomas Zhou, Eric Du, Thiago Missfeldt Sanches, Khai-Minh Nguyen, Mika Okamoto, Abraham A Palmer, Rick Friedman

## Abstract

Age-related hearing impairment is the most common cause of hearing loss and is one of the most prevalent conditions affecting the elderly globally. It is influenced by a combination of environmental and genetic factors. The mouse and human inner ears are functionally and genetically homologous. Investigating the genetic basis of age-related hearing loss (ARHL) in an outbred mouse model may lead to a better understanding of the molecular mechanisms of this condition. We used Carworth Farms White (CFW) outbred mice, because they are genetically diverse and exhibit variation in the onset and severity of ARHL. The goal of this study was to identify genetic loci involved in regulating ARHL. Hearing at a range of frequencies was measured using Auditory Brainstem Response (ABR) thresholds in 946 male and female CFW mice at the age of 1, 6, and 10 months.

We obtained genotypes at 4.18 million single nucleotide polymorphisms (SNP) using low-coverage (mean coverage 0.27x) whole-genome sequencing followed by imputation using STITCH. To determine the accuracy of the genotypes we sequenced 8 samples at >30x coverage and used calls from those samples to estimate the discordance rate, which was 0.45%. We performed genetic analysis for the ABR thresholds for each frequency at each age, and for the time of onset of deafness for each frequency. The SNP heritability ranged from 0 to 42% for different traits. Genome-wide association analysis identified several regions associated with ARHL that contained potential candidate genes, including *Dnah11*, *Rapgef5*, *Cpne4*, *Prkag2*, and *Nek11*. We confirmed, using functional study, that Prkag2 deficiency causes age-related hearing loss at high frequency in mice; this makes *Prkag2* a candidate gene for further studies. This work helps to identify genetic risk factors for ARHL and to define novel therapeutic targets for the treatment and prevention of ARHL.

## Introduction

Age-related hearing loss (ARHL) is the most common cause of hearing loss and is one of the most prevalent conditions affecting the elderly globally. Twin and family studies reveal 25-75% of risk for ARHL is due to heredity (Momi et al. 2015). There is very little existing information about the genes and pathways responsible for ARHL in humans and mice despite the evidence from our lab and others supporting its heritabiltiy and polygenic architecture (Fransen et al. 2015). Estimates suggest that approximately two-thirds of people over the age of 70 in the United States experience ARHL (Bainbridge and Wallhagen, 2014). ARHL has been shown to be independently associated with cognitive decline, dementia, depression, and loneliness and results in an estimated annual economic burden of over $3 billion in medical expenditures (Deal et al. 2017; Deal et al. 2018; Lin and Albert, 2014). Although the use of hearing aids and/or cochlear implants has been shown to improve many of these associated conditions, ARHL remains significantly undertreated and to date, there are no targeted therapies (Deal et al. 2018).

Greater than 100 genes have been identified in association with monogenic, non-age-related deafness. However, a substantial fraction of patients with ARHL have no identifiable mutation in any known hearing loss gene, suggesting that a significant fraction of hearing loss is due to a combination of environmental causes and unidentified monogenic or polygenic causes (Bowl and Dawson, 2018). There is ample evidence that the anatomical, cellular, and molecular properties of the mouse and human inner ear are highly homologous. All mammalian inner ear development begins with a thickening of the ectoderm (otic placode) (Bok et al. 2007).

The placode then invaginates to form the otocyst. Within the otocyst Sox2-positive epithelial prosensory patches are specified, one of which gives rise to the cochlea. As the cochlear duct extends, there is a wave of differentiation within and surrounding the duct. Ultimately, the cochlea consists of three fluid-filled spaces, the scala vestibuli, scala media, and scala tympani. The sensory and supporting cells exist within the organ of Corti and reside within the scala media. In a recent manuscript describing mouse and human inner ear development at the single-cell level, the authors concluded: “Our analysis revealed remarkable similarity between human and mouse cell cochlear subpopulations. We observed a remarkably similar pattern of key markers of distinct subpopulations in the developing human cochlea to the developing mouse cochlea. Thus, we believe that despite the size and timing differences of cochlear development between the mouse and human, the mouse is likely a very good model of human cochlear development” (Yu et al. 2019).

Genome-wide association studies (GWAS) of hearing traits in humans, including ARHL, have identified a few genome-wide significant risk loci, but many suffer from a lack of sufficient power (Friedman et al. 2009; Girotto et al. 2011; Hoffmann et al. 2016; Van Laer et al. 2010; Vuckovic et al. 2015; Praveen et al. 2022). Our laboratory previosuly performed the first human GWAS for ARHL in which we identified a genome-wide significant risk locus within intron 2 of *GRM7* (Friedman et al. 2009). *GRM7* was subsequently implicated in ARHL by other genetic studies as well (Newman et al. 2012; Van Laer et al. 2010). Recently, two ARHL GWAS used data from the UK Biobank (UKBB), which includes genotype and questionnaire (no formal audiograms) data from more than 330,000 individuals, identified several genome-wide significant associations, some of which were in or near genes that cause Mendelian deafness (Kalra et al. 2020; Wells et al. 2019). Notably, all of the candidate genes described were significantly enriched with mouse phenotype ontologies, mostly related to mouse inner ear abnormalities and abnormal auditory brainstem response (ABR) with the authors concluding “this finding demonstrates the shared genetic pathology in mouse and human auditory systems, supporting the use of mouse models to study human auditory function” (Wells et al. 2019). Further support comes from our recent work identifying altered expression levels of *Fhod3*, on mouse chromosome 18, which results in reduced actin content in the cuticular plate, loss of the third-row stereocilia in the cochlear base, and progressive high frequency hearing loss (Boussaty et al. 2023). A recent analysis of hearing loss diagnoses in the Million Veteran Program (MVP) via GWAS revealed several loci containing genes associated with stereociliary structure and function in much the same way as *Fhod3* (De Angelis et al. 2023).

Mouse GWAS have several advantages: the environment can be more carefully controlled, a greater proportion of the heritability can be captured, and findings can be followed up using experimental manipulations. The Knockout Mouse Project/International Mouse Phenotyping Consortium (KOMP-IMPC) has identified 62 novel genes involved in early onset hearing loss by testing Auditory Brainstem Response thresholds in 14 week-old mice (http://www.mousephenotype.org) (Bowl et al. 2017). While this valuable resource may assist with modeling candidate genes discovered in this work, and we see some overlap, these are young mice for ARHL (14 weeks) and are null mutations and therefore the study was biased to Mendelian forms of congenital (rather than age-related) deafness. Furthermore, KOMP-IMPC will miss embryonic lethal genes, even when modest decreases in those genes’ function may produce viable animals with hearing-related phenotypes.

This paper presents the genetic analysis of the hearing function of the aging outbred Carworth Farms White (CFW) mice measured through the auditory brainstem response (ABR) thresholds (Du et al. 2022).

Commercially available CFW outbred mice (Lynch, 1969) have reduced linkage disequilibrium (Parker et al. 2016) and provide fine-scale mapping resolution that is better than panels of inbred strains or other commercially available oubred mice (Bowl et al. 2017, Yalcin et al. 2010). The heterogeneity of ABR thresholds in genetically diverse CFW mice provides an opportunity to study the genetic landscape of ARHL. We are working to continue to accelerate the pace of discovery of polygenic loci and pathways for ARHL through a novel “forward genetics” approach and our initial GWAS results are the subject of this manuscript.

## Methods

### Animals

All procedures were performed in accordance with guidelines from the National Institutes of Health and the Association for the Assessment and Accreditation of Laboratory Animal Care and approved by the Institutional Care and Use Committee at the University of California San Diego. The detailed procedures are reported in (Du et al. 2022). Briefly, the animals were obtained from the Crl:CFW(SW)-US_P08 (CFW) stock of outbred mice maintained by Charles River Laboratories (Portage, MI). The mice arrived at 3 weeks of age.

They were tested at 1, 6, and 10 months (**Table 1**, **Fig. 1A**). We requested that only one mouse from one litter was shipped, to avoid using siblings, which reduces the power of GWAS. Subsequent genetic analysis demonstrated that about 252 out of 946 mice used in this work were siblings (**Supplemental Fig. 1**).

**Figure 1.**
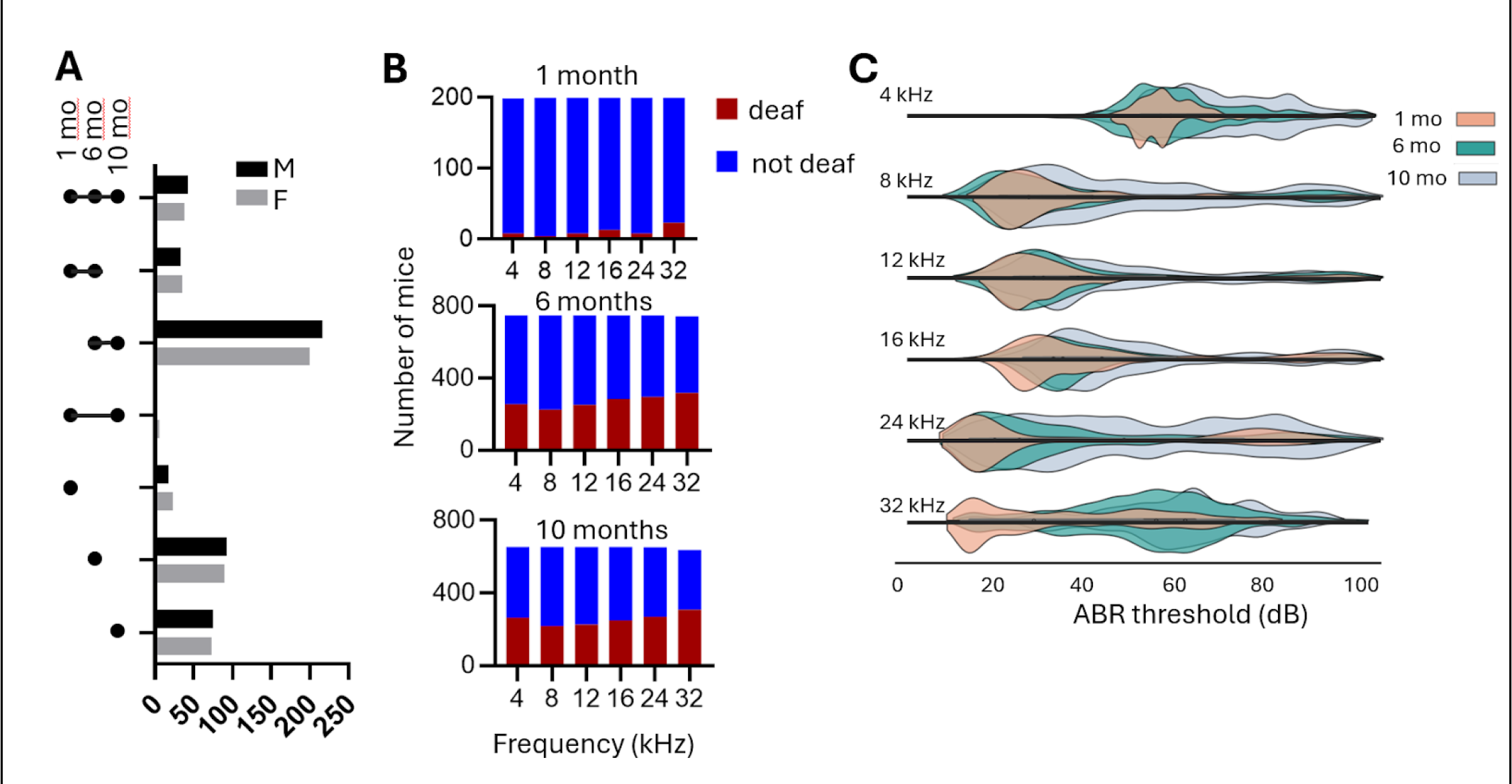
Mice and phenotypes used for genetic analysis. A. . Mice used in the experiment. Not all animals have measurements at all ages, see Table Cohort for the summary. **B** .The number of CFW mice that become deaf and retain hearing at different frequencies at three time points. **C** . Ridgeplot of the ABR thresholds in CFW mice measured at three ages at different frequencies. Females are shown above the X-axis, and males are shown below the X-axis for each frequency. The ABR>100 is considered “deaf” and not included in this plot.

**Table 1.** Demographic description of subjects.

| Age group | Males |  |  | Females |  |  | Total |
| --- | --- | --- | --- | --- | --- | --- | --- |
|  | N | Age mean, days | Age st. dev, days | N | Age mean, days | Age st. dev, days | N |
| 1 month | 97 | 46.5 | 9.6 | 102 | 47.2 | 9.8 | 199 |
| 6 month | 385 | 197.8 | 14.1 | 363 | 194.5 | 11.9 | 748 |
| 10 month | 337 | 313.4 | 13.1 | 317 | 315 | 13.2 | 654 |

Animals were housed 3 per cage with a low lever of ambient noise, on a standard 12:12 h light–dark cycle, standard laboratory chow, and water *ad libitum*. Phenotyping occurred during the light phase. Spleens were harvested after the mice were sacrificed, and used as a source of DNA for genotyping.

### Auditory brainstem response testing

The ABR thresholds were measured at three time points: 5-8 weeks (denoted as “1 month” in this paper), 6 months, and 10 months of age, as described earlier (Du et al. 2022). Briefly, the mice were anesthetized using ketamine (80 mg/kg) and xylazine (16 mg/kg) intraperitoneal injection. All hearing tests were performed in a soundproof acoustic chamber. Stimuli were provided by a custom acoustic system consisting of two miniature speakers with sound pressure measured by a condenser microphone.Auditory signals were presented as tone pips with a rise and a fall time of 0.5 msec and a total duration of 5 msec at the frequencies 4 kHz, 8 kHz, 12 kHz, 16 kHz, 24 kHz, and 32 kHz. These tone pips started at 20 dB and then increased in 5 dB increments up to 100 dB SPL and were presented at a rate of 30/second. The responses were recorded and then filtered with a 0.3 to 3 kHz pass-band. 350 waveforms were averaged for each stimulus intensity. Hearing thresholds were determined by visual inspection; if no wave form was detected at 100 dB SPL, the hearing threshold was recorded as “no response” indicating that a mouse is deaf at this frequency.

### Genotyping

DNA was extracted from mouse spleen tissue using DNAdvance kit (Beckman Coulter). Multiplexed sequencing libraries were prepared using the Twist 96-Plex Library Prep kit (TWIST Bioscience), and then sequenced on a NovaSeq 6000 or NovaSeq X (Illumina). An average of ∼3.2 million reads per sample were obtained (paired end, 150 bp). The reads were aligned to the mouse reference genome GRCm38 (GCA_000001635.2). To generate genotypes at single nucleotide polymorphisms (SNPs) we used STITCH software (Davies et al. 2016) without reference panel, with the “niterations” parameter set to 40 and a position file as described below, to create a reference panel; then we ran STITCH again with the “niterations” parameter set to 1, using with the above result as a reference panel, filtered resulted genotypes by INFO score > 0.9, and then performed imputation using BEAGLE software (Browning and Browning, 2016) . To construct the position file we used low-coverage sequencing data from earlier generations of CFW mice (Zou et al. 2022; Davis et al. 2016) and from 8 CFW individuals from the current study sequenced at 30x (see below). The SNPs on X, Y, and MT chromosomes were not called, in part because only autosomes were available in the (Zou et al. 2022) dataset. After genotypes were called with STITCH, the following SNPs were removed from the analysis: (1) monomorphic SNPs, since they are not useful for further genetic analysis, (2) SNPs that violate Hardy-Weinberg equilibrium (HWE) with -log_10_(p) > 7, where p is the p-value of the HWE test at a SNP, and (3) SNPs that have genotype missingness rate > 0.1 based on the results generated by STITCH. The filtered dataset contained ∼4.18 million SNPs on 19 autosomes (**Fig. 2**). The data is available from UCSD Library (doi.org/10.6075/j0h13263). The animals with SNP missingness rate > 0.2 were removed from the analysis. The final dataset for genetic analysis consisted of 946 CFW mice. Eight of those same CFW mice were also sequenced at >30x coverage. Genotypes for the >30x data were called by GATK and filtered using bcftools to exclude the following SNPs: (1) not biallelic, (2) QUAL < 20, (3) GQ < 20, and (4) genotype missing rate >= 0.15. The 8 deeply sequenced samples were used as a “truth set” to determine the accuracy of the genotyping and estimate the error rate, which was 0.45%.

**Figure 2.**
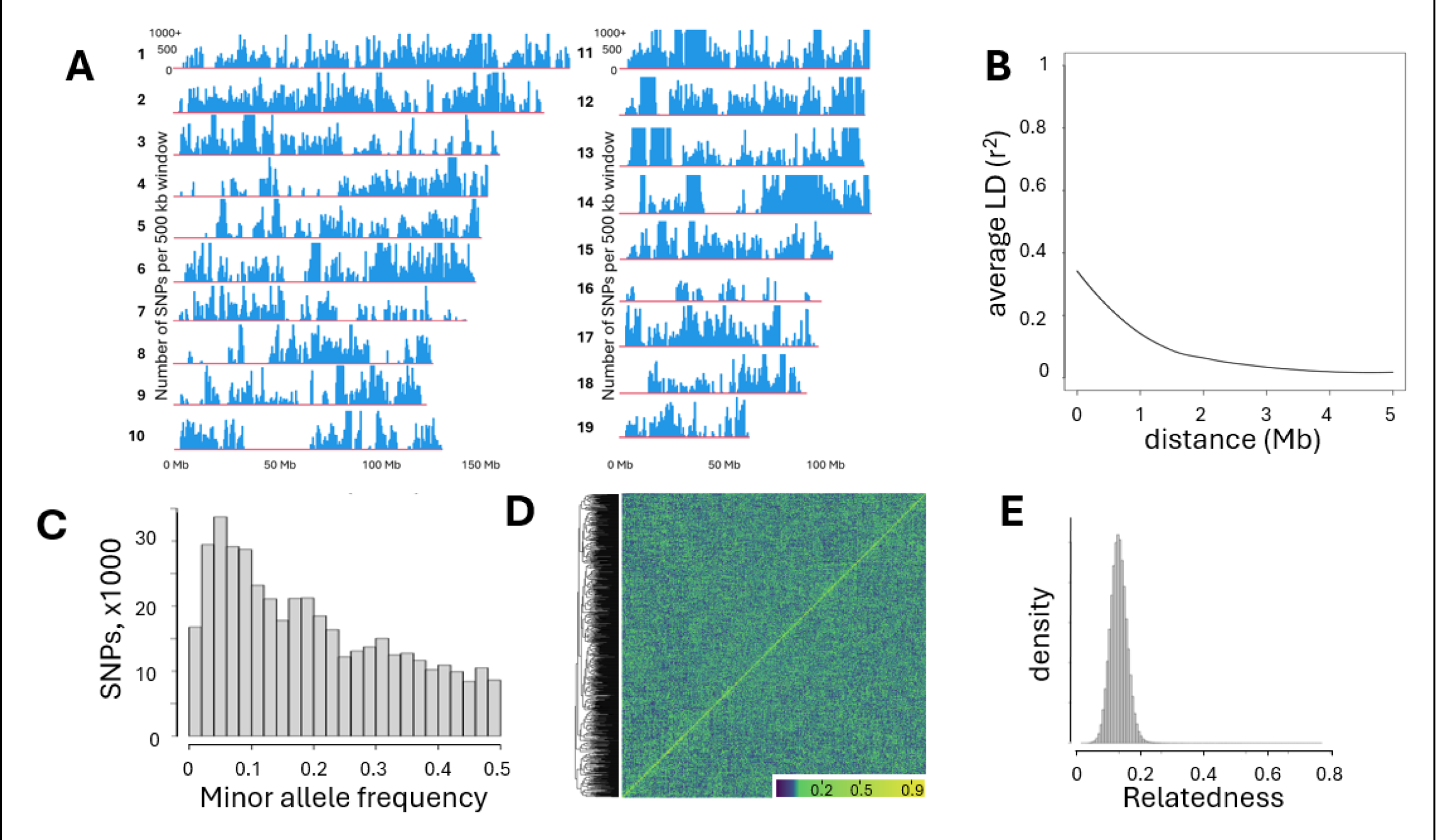
Genetic architecture of CFW mice. A. Density of SNPs called in autosomes 1- 19; bin size is 500 kb. **B.** LD decay in CFW mice used in this study. **C.** Histogram of SNP number with MAF>0. **D.** Heatmap of the genetic relatedness matrix (GRM) of the genotypes of CFW mice in this study demonstrates an absence of a noticeable genetic structure; samples clustered by genetic relatedness. **E** . Relatedness calculated as PiHat among all pairs of CFW mice in this study shows that most of the mice used for the study are not closely related.

The extent of LD (r^2^) decay rates in CFW mice was estimated as follows: bcftools was used to create the set of SNPs pruned to remove sites with r^2^ > 0.95 within 5Mb, and then vcftools was used to calculate LD metric r^2^ for each pair of SNPs that are within 5 Mb. The relationship curve between physical distance and r^2^ was fitted by LOESS (locally estimated scatterplot smoothing).

### Genetic analysis

For genetic analysis, each quantitative trait was quantile-normalized. Sex was used as a covariate if it explained more than 2% of the variance. There were 2 traits where sex was used as a covarite: ABR threshold at 4 khz and at 12 kHz at 1 month of age; the sez explained 3% of variance for these traits. The SNP heritability was estimated using GCTA-GREML (Yang et al. 2010). Genetic correlations between traits were computed through bivariate GREML analysis performed with GCTA (Lee et al. 2012). GWAS analysis was performed using a linear mixed model, as implemented in GCTA (Yang et al. 2011), with the genetic relatedness matrix (GRM) used to account for the complex family relationships within the CFW population, and the Leave One Chromosome Out (LOCO) method to avoid proximal contamination (Cheng et al. 2013; Gonzales et al. 2018). LOCO, which was coined by (Yang et al. 2014) and first proposed by (Cheng et al. 2013), is a computationally efficient strategy to address the concern of “proximal contamination” (Listgarten et al. 2012) that can reduce the statistical power of GWAS. To control for the type I error, the significance threshold was estimated by a permutation test (Cheng and Palmer, 2013). We used permutation to establish the genome-wide significance thresholds. The genome-wide thresholds for -log10(p) at the significance levels 0.05 and 0.10 were 5.58 and 5.0 respectively. Quantitative trait loci (QTL) were determined by at least one SNP that exceeded the permutation-derived threshold of −log_10_(p) > 5.0, which was supported by a second SNP within 0.5 Mb of this SNP that had a p-value that was within 2 − log_10_(p) units of the most significant SNP. Regional association plots were generated using LocusZoom software (Pruim et al. 2010).

## Results

### Age-related decrease in auditory brainstem response

ABR thresholds were measured in 946 outbred CFW mice (**Fig. 1A)**. As expected, an increasing proportion of mice became deaf with increased age (**Fig. 1B)**, and in those that retained hearing the ABR thresholds increased with age (**Fig. 1C**). A detailed description of the hearing loss patterns in CFW mice was published by (Du et al. 2022) for a subset of the mice that were used for the genetic analysis; however, the description and conclusions can be applied to the cohort used in this work. We performed two-way ANOVA to determine where there is a difference between males and females.

The main effect of sex was statistically significant (p < 0.001, F = 28.16, df = 1). The interaction effect between sex and time point was also significant (p = 0.015, F = 4.19, df = 2).

For the genetic analysis, we considered two types of phenotypes: “deaf vs. not deaf at each frequency” and “ABR threshold at each frequency”.

### Genetic architecture of the CFW mice

Polymorphic SNPs were unevenly distributed across the autosomes; we identified several large regions with few or no polymorphic SNPs, which is consistent with prior studied using CFW mice (Parker et al. 2016) and likely reflects both a true lack of diversity and regions that may be highly repetitive or otherwise difficult to genotype (**Fig. 2A**). LD decay in this population supports its suitability for high-resolution mapping (**Fig. 2B**). The distribution of minor allele frequencies (MAF) of SNPs shows that 85.6% of non-monomorphic SNPs had MAF > 0.05, which is consistent with the history of CFW mice (**Fig. 2C**). The genetic structure of the population that can be present due to breeding schema in vendor’s facilities can skew the genetic analysis (Gileta et al. 2022), therefore we checked that the CFW population used in this study was genetically homogeneous. A heatmap of the genetic relatedness matrix (GRM) demonstrates an absence of a noticeable genetic structure (**Fig. 2D**). Relatedness calculated as π-hat among all pairs of CFW mice in this study shows that most of the mice used for the study are not closely related (**Fig. 2E**).

<u>Heritability</u> estimates for hearing thresholds and hearing loss traits range between 0 and 42% (**Table 2**). The SNP heritability is expected to be lower than the heritability estimated in twin studies or by using inbred strains. In this dataset, the heritability was moderate across the different measures of hearing.

**Table 2.** Heritability of traits.

| Table 2. Heritability of traits |  |  |  |  |
| --- | --- | --- | --- | --- |
| Trait | N | SNP heritability | SNP heritability SEM | p-value |
| deaf_06mo_04khz | 748 | 0.106 | 0.055 | 0.02 |
| deaf_06mo_08khz | 748 | 0.054 | 0.052 | 0.157 |
| deaf_06mo_12khz | 748 | 0.1 | 0.056 | 0.034 |
| deaf_06mo_16khz | 748 | 0.148 | 0.059 | 0.003 |
| deaf_06mo_24khz | 748 | 0.19 | 0.06 | 0 |
| deaf_06mo_32khz | 745 | 0.187 | 0.059 | 0 |
| deaf_10mo_04khz | 654 | 0.132 | 0.063 | 0.01 |
| deaf_10mo_08khz | 654 | 0 | 0.048 | 0.5 |
| deaf_10mo_12khz | 654 | 0.06 | 0.054 | 0.11 |
| deaf_10mo_16khz | 653 | 0.047 | 0.055 | 0.18 |
| deaf_10mo_24khz | 651 | 0.032 | 0.053 | 0.263 |
| deaf_10mo_32khz | 636 | 0.033 | 0.055 | 0.264 |
| thr_01mo_04khz | 190 | 0 | 0.154 | 0.5 |
| thr_01mo_08khz | 195 | 0.346 | 0.166 | 0.014 |
| thr_01mo_12khz | 191 | 0.212 | 0.171 | 0.094 |
| thr_01mo_16khz | 186 | 0.18 | 0.198 | 0.222 |
| thr_01mo_24khz | 191 | 0.417 | 0.169 | 0.005 |
| thr_01mo_32khz | 176 | 0.22 | 0.193 | 0.142 |
| thr_06mo_04khz | 493 | 0.086 | 0.074 | 0.109 |
| thr_06mo_08khz | 523 | 0.208 | 0.076 | 0.001 |
| thr_06mo_12khz | 497 | 0.109 | 0.072 | 0.044 |
| thr_06mo_16khz | 464 | 0.123 | 0.082 | 0.048 |
| thr_06mo_24khz | 453 | 0.035 | 0.074 | 0.312 |
| thr_06mo_32khz | 427 | 0.2 | 0.099 | 0.023 |
| thr_10mo_04khz | 390 | 0.089 | 0.097 | 0.18 |
| thr_10mo_08khz | 437 | 0.226 | 0.096 | 0.006 |
| thr_10mo_12khz | 427 | 0.106 | 0.094 | 0.144 |
| thr_10mo_16khz | 405 | 0.176 | 0.097 | 0.024 |
| thr_10mo_24khz | 382 | 0.23 | 0.108 | 0.014 |
| thr_10mo_32khz | 329 | 0.188 | 0.126 | 0.078 |

### Genetic correlations

To examine the genetic relatedness among thresholds and deafness, genetic correlations were computed (**Figure 3**). Correlations were not calculated for the “deaf at 1 month” traits because of an imbalanced number of deaf mice and a small sample size. The deafness at 6 month for any frequency had strong genetic correlates with deafness at 6 month for all frequencies. Similarly, ABR thresholds at 6 month at any frequency had strong genetic correlation with ABR thresholds at 6 month at other frequencies. The ABR thresholds at 1 month tend to have negative correlation with ABR thresholds at 10 month.

**Figure 3.**
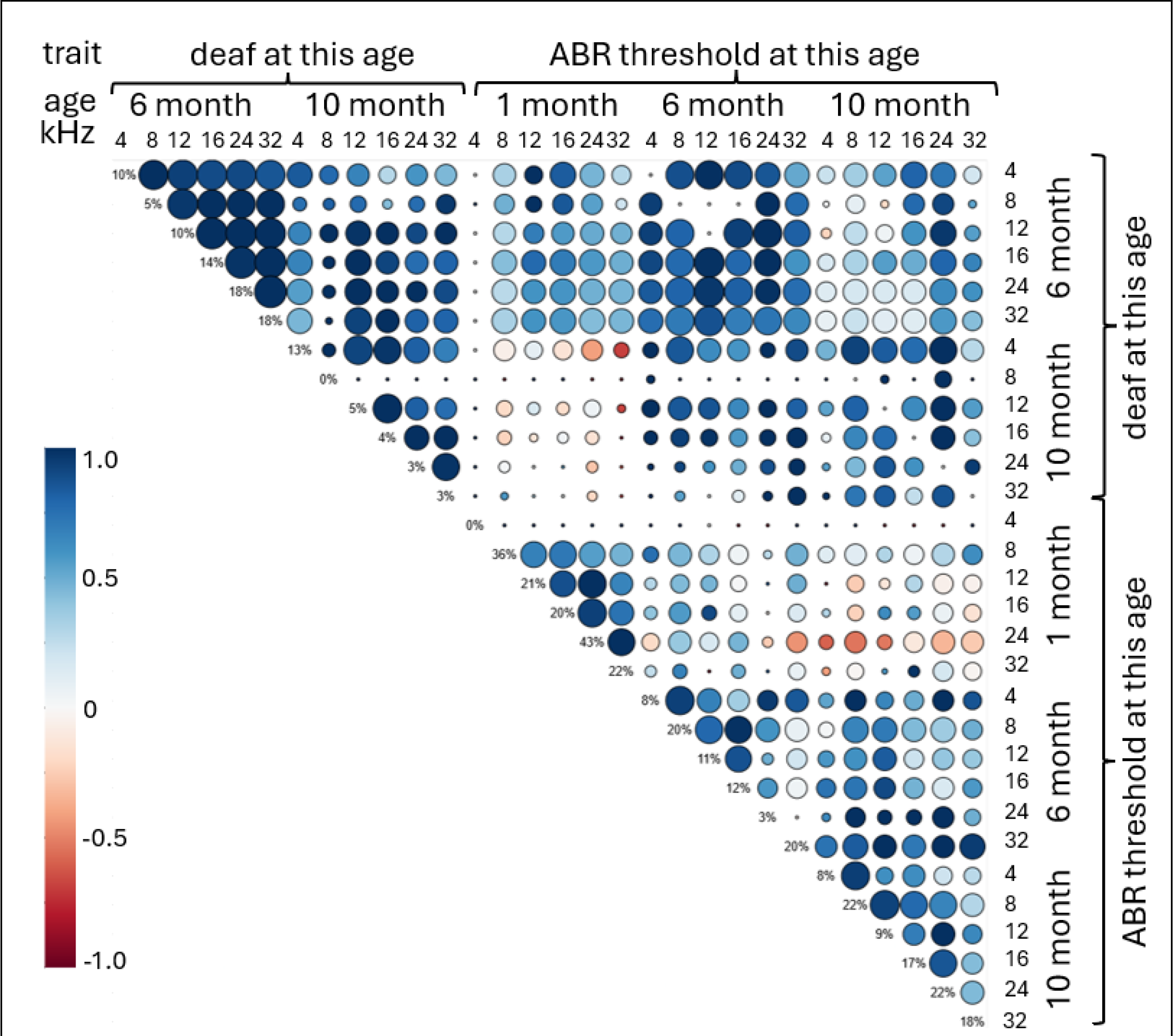
Genetic correlation for the hearing traits measured at three time points. The color of the circle indicates correlation. The size of the circle shows significance of the correlation. The values on the diagonal show heritability for each trait.

### GWAS results

GWAS was performed to identify loci that were significantly associated with hearing thresholds and with deafness for each threshold, at each timepoint. We identified 10 QTLs for 7 traits. The list of QTLs with effect sizes and top SNP frequencies is shown in **Table 3**. The chromosomal locations for identified regions of interest are shown as a porcupine plot (**Figure 4**). The full genetic report is available in **Supplemental Materials**.

**Figure 4.**
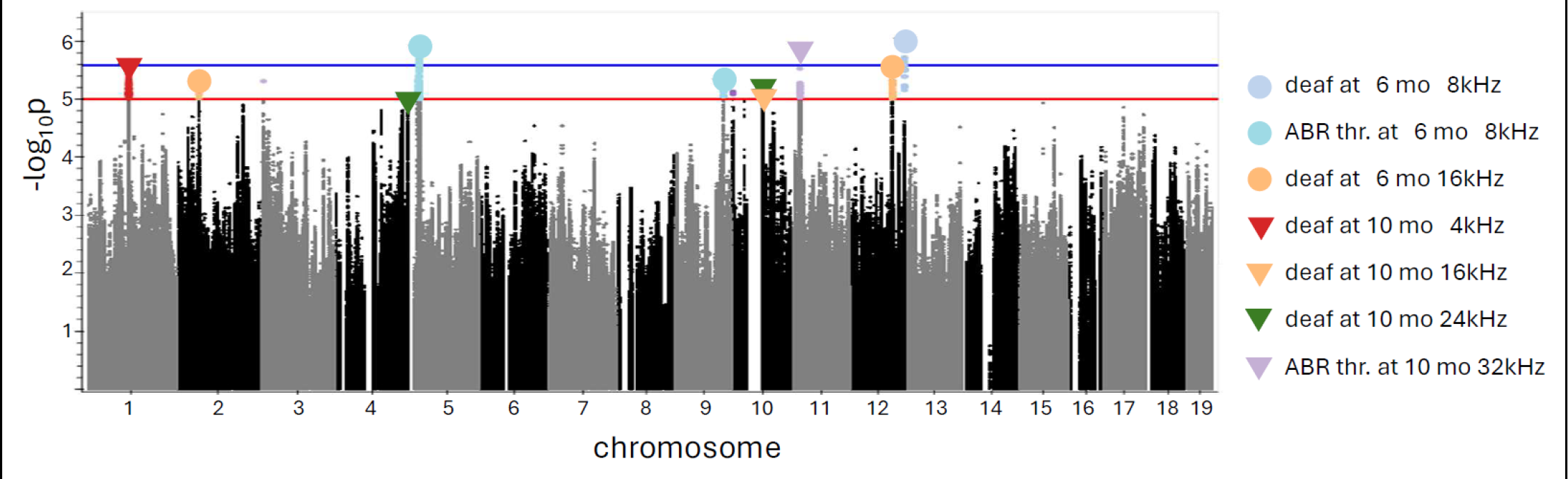
Porcupine plot for all measured ARHL traits. The red line indicated a threshold for genome-wide alpha of < 0.10 (-log10(p) > 5.0); the blue line indicated a threshold for genome-wide alpha of < 0.05 (-log10(p) > 5.58). Triangles indicate top SNPs, with colors showing a specific trait.

**Table 3.** Description of QTLs.

| Trait | Top SNP | Top SNP allele | Effect size | Effect size SE | $-\log_{10}(p)$ for the top SNP | Number of genes |
| --- | --- | --- | --- | --- | --- | --- |

|  |  | frequency |  |  |  |  |
| --- | --- | --- | --- | --- | --- | --- |
| deaf at 06 mo 08 khz | chr12:117,951,339 | 0.140 | 0.181 | 0.037 | 5.99 | 4 |
| deaf at 06 mo 16 khz | chr2:49,860,512 | 0.183 | -0.153 | 0.034 | 5.24 | 2 |
| deaf at 06 mo 16 khz | chr12:91,900,313 | 0.951 | -0.274 | 0.059 | 5.52 | 2 |
| deaf at 10 mo 04 khz | chr1:95,110,035 | 0.844 | 0.190 | 0.040 | 5.56 | 0 |
| deaf at 10 mo 16 khz | chr10:67,242,708 | 0.315 | 0.141 | 0.031 | 5.24 | 5 |
| deaf at 10 mo 24 khz | chr4:155,998,026 | 0.051 | 0.283 | 0.064 | 5.00 | 6 |
| deaf at 10 mo 24 khz | chr10:67,229,071 | 0.315 | 0.142 | 0.032 | 5.11 | 5 |
| ABR threshold at 06 mo 08 khz | chr5:24,688,891 | 0.146 | 0.434 | 0.090 | 5.87 | 35 |
| ABR threshold at 06 mo 08 khz | chr9:105,380,885 | 0.057 | 0.605 | 0.133 | 5.29 | 5 |
| ABR threshold at 10 mo 32 khz | chr11:16,309,495 | 0.132 | -0.578 | 0.119 | 5.90 | 2 |

Due to the high resolution of genetic mapping in the CFW population most QTLs contained only a few genes. Each QTL interval was examined at the Mouse Genome Informatics portal (MGI) (Blake et al. 2021) which aggregates data on previously reported QTLs and mutant phenotypes, as well as gene expression. To identify candidate genes within each QTL, we considered several criteria: whether the gene was located within an interval that contains SNP in a high LD with the top SNP (r^2^ > 0.8), the presence of moderate or high impact variants located within the gene, as predicted by SnpEff (Cingolani et al., 2012), and the expression in the tissue of interest, cochlea, in the publicly available datasets that are available in the MGI database and the gEAR portal (umgear.org). In addition, the dataset from (Boussaty et al. 2023) was obtained from 48 10-month old CFW mice with and without hearing loss; 45 of the mice from (Boussaty et al. 2023) are included in the current study. Five of the genes that were found in the QTL regions identified in this study were also detected in (Boussaty et al. 2023), and are discussed below.

We detected three QTLs for elevated thresholds at 6 months (genetic report in **Supplemental Materials**).

<u>QTL on chromosome 12 at around 118 Mb</u> (**Figure 5A).** No hearing-related mutations or QTLs have been reported for this locus in any organisms, this work is the first report. This QTL is associated with being deaf at 6 months at 4 kHz, and also shows a trend towards an association with elevated thresholds at 6mo at 12kHz (-log10(p) = 4.6).There are 4 genes in this locus: *Dnah11* (dynein, axonemal, heavy chain 11), Cdca7l (cell division cycle associated 7 like), *Sp4* (trans-acting transcription factor 4), *Rapgef5* (Rap guanine nucleotide exchange factor 5).

**Figure 5.**
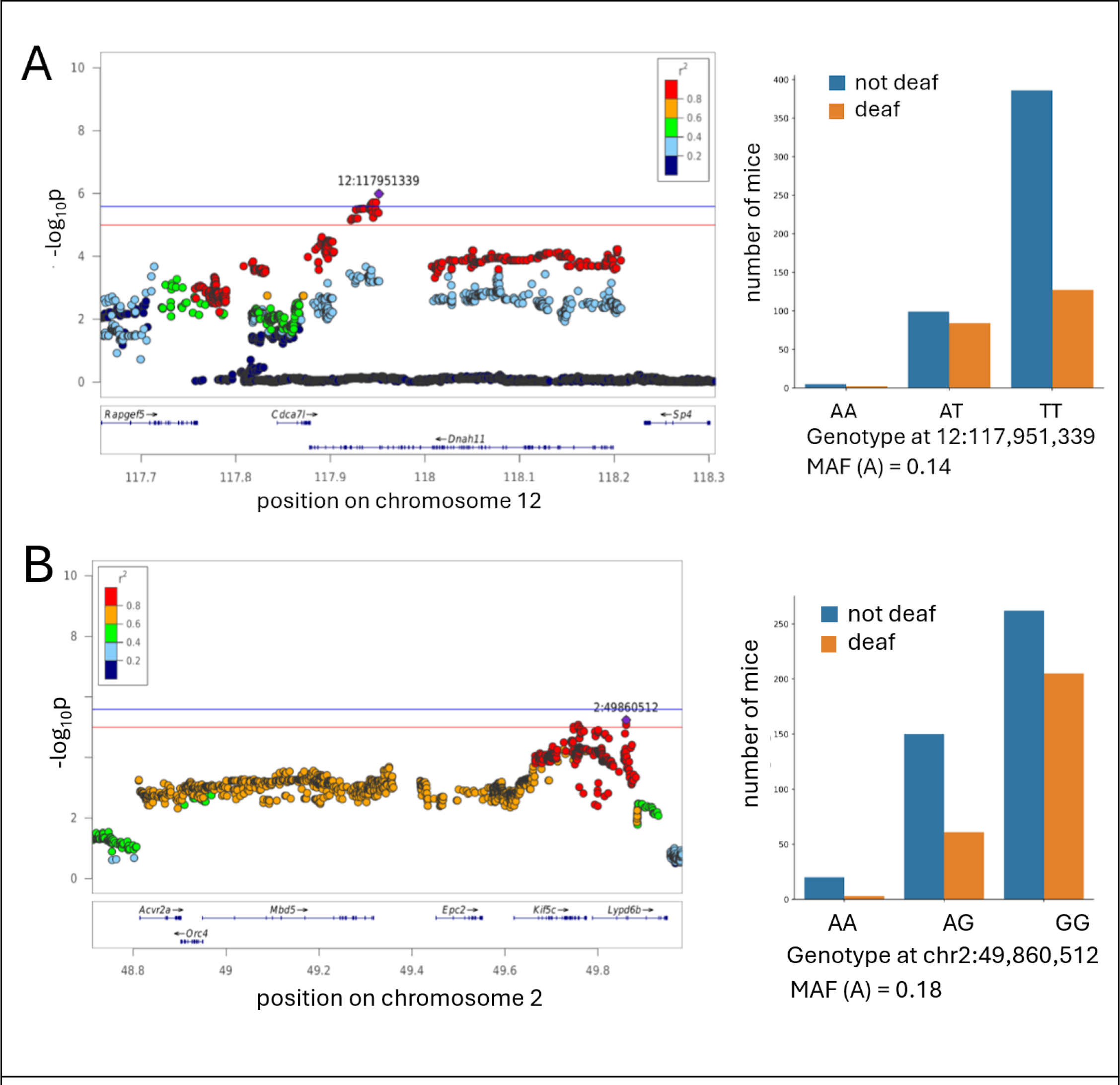
QTLs for the traits of being deaf at 6 months. The x-axis shows the position on a chromosome (in Mb); the y-axis shows the significance of the association (-log10 p-value). The individual points represent SNPs. The SNP with the most significnat p-value (“top SNP”) is highlighted in purple. The colors represent the LD between the topSNP and the other SNP. The red line indicates the threshold for genome-wide alpha of < 0.10 (-log10(p) > 5.0); the blue line indicates the threshold for genome-wide alpha of < 0.05 (-log10(p) > 5.58). The effect plots for the top SNP are shown on the right, minor allele frequency (MAF) indicated.

*Dnah11* has no previously reported connection to hearing/vestibular phenotypes in mice, according to MGI database. In humans, mutations in this gene cause diseases related to the dysfunction of the cilia during embryologic development, such as situs inversus (abnormal distribution of the major visceral organs within the chest and abdomen). Dnah11 is expressed in hair cells in 10-month old CFW mice (Boussaty et al. 2023).

Other genes that encode dynein heavy chains are known to cause hearing dysfunction: *DNAH2* is predicted to cause ARHL in humans, as determined by presence of a rare variant tha affects protein function in a cohort of hearing loss patients (Lewis et al. 2018); another related gene *Dync1li1* is required for the survival cochlear hair cells in mice (Zhang et al. 2022). In CFW mice *Dnah11* has two missense mutations, both in linkage disequilibrium with the top SNP: 12:118,190,825 (c719A>G; Glu240Gly, r^2^ with the top SNP 0.982) and 12:118,198,712 (c121C>T; Arg41Cys, r^2^ with the top SNP 0.972).

*Rapgef5* (Rap guanine nucleotide exchange factor 5) has not been previously associated with hearing/vestibular phenotypes in any species. An indel mutation in RAPGEF5 causes epilepsy in dogs (Belanger et al. 2022). The function of RapGEFs can vary depending on the specific isoform and the cellular context, they regulate cell adhesion, cytoskeletal dynamics, and tissue morphogenesis; these processes are important for the development and maintenance of the structures of the inner ear, making *Rapgef5* a plausible novel candidate gene. In 10-month old CFW mice, this gene was found to be expressed in endothelial cells, border cells and pillar cells (Boussaty et al. 2023).

<u>QTL on chromosome 2 at 49.7 Mb</u> (**Figure 5B**). This locus was associated with being deaf at 6 months at 16 kHz, and also trending towards associations with hearing loss at 6 months for all other tested frequencies, although these associations do not reach the significance threshold (-log10(p) range from 4.01 to 4.55). There are two genes in this locus: *Kif5c* (kinesin family member 5C) and *Lypd6b* (LY6/PLAUR domain containing 6B).

*Kif5c* (kinesin family member 5C) encodes a protein which has microtubule motor activity and is located in the related cellular components, including ciliary rootlet. Although we are not aware of any previously reports that *Kif5c* is involved in hearing in any species, other kinesin motor proteins are known to be associated with hearing loss. For example, *Klc2* (kinesin light chain 2) knock out mice have early hearing loss at low frequencies, and KLC2 binds KIF5C in the mouse cochlea, as shown in co-immunoprecipitation experiments (Fu et al. 2021). This gene was not detected in 10-month old CFW mice (Boussaty et al. 2023), but is expressed in inner hair cells in young mice (Liu et al. 2018).

<u>QTL on chromosome 12 at 92 Mb.</u> (genetic report in **Supplemental Materials**). The most strongly associated SNP for this QTL also showed a trend towards an association with hearing loss at 6 months for 12 kHz and 24 kHz (-log10(p)=4.4 and 5.02). This locus contains genes *Ston2* (stonin 2) and *Sel1l* (sel-1 suppressor of lin-12-like). These two genes are not known to be associated with hearing loss and were not expressed in 10-month old CFW mice (Boussaty et al. 2023), however *Sel1l* was expressed in cochlea cells in several of the gEAR datasets (Sun et al. 2023; Su et al. 2019).

We detected three QTLs for being deaf at 10 months (genetic report in **Supplemental Materials**).

<u>QTL on chromosome 1 at 95.2 Mb</u> (genetic report in **Supplemental Materials**). This locus is associated with hearing loss at 10 months at 4 kHz, and also showed a trend towards an association with being deaf at 10 month for the 16 kHz and 32 kHz frequencies (-log10(p)= 4.26 and 4.13 correspondingly). This locus does not contain any known genes, but may contain unknown genes or transcripts or regulatory sequences that influences the expression of genes outside the associated region. As far as we know, this locus has not been previously associated with hearing loss in any species.

<u>QTL on chromosome 10 at 67 Mb</u> (**Figure 6**). This chromosomal region contains QTLs for hearing loss at 10 months for both 16 kHz and 24 kHz. . It contains 3 genes: *Reep3* (receptor accessory protein 3), *Jmjd1c* (jumonji domain containing 1C), and the predicted gene *Gm31763*.

**Figure 6.**
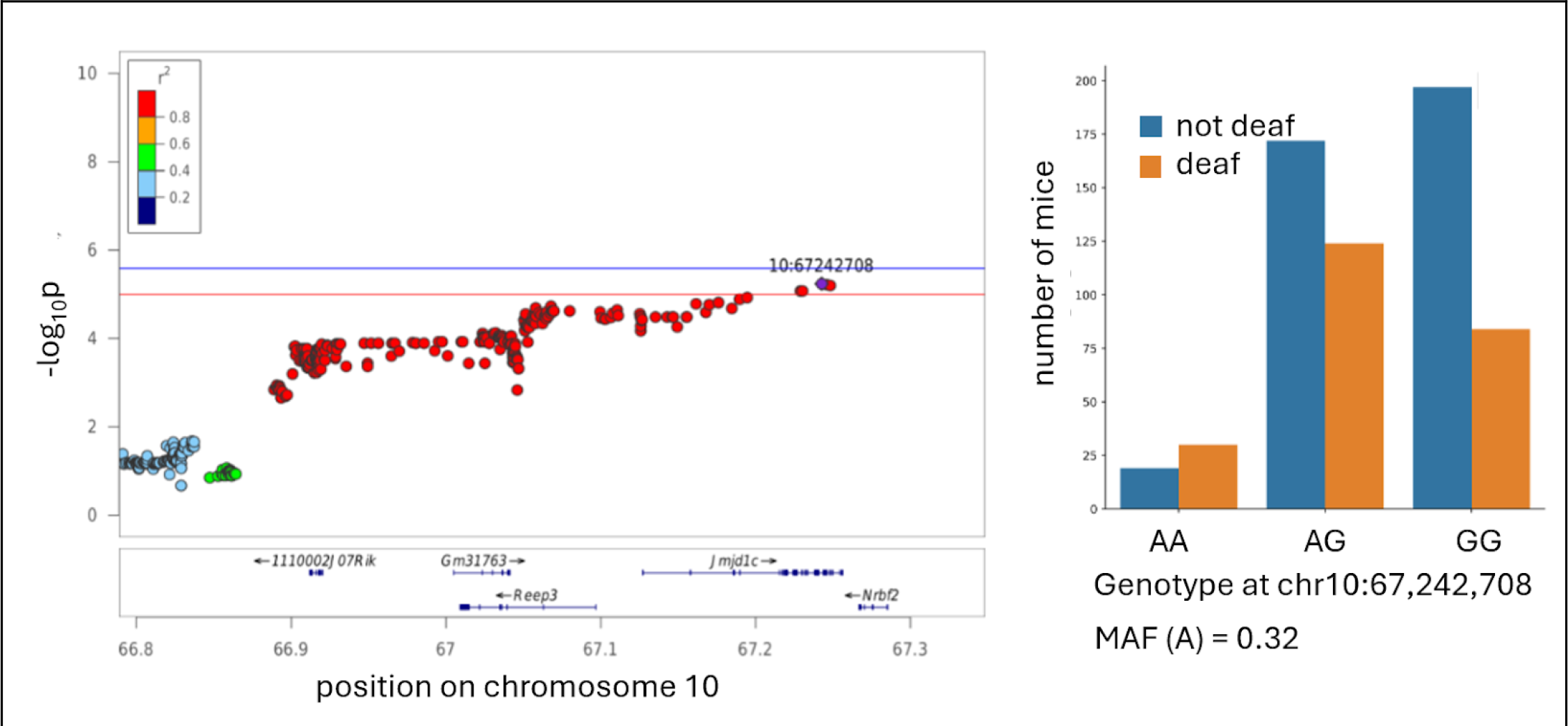
QTL for the trait of being deaf at 10 month. The regional association plots are shown on the left. The x-axis shows the position on a chromosome (in Mb); the y-axis shows the significance of the association (-log10 p-value). The individual points represent SNPs. The SNP with the lowest p-value (“top SNP”) is highlighted in purple. The colors represent the correlation between the topSNP and the other SNP. The red line indicated a threshold for genome-wide alpha of < 0.10 (-log10(p) > 5.0); the blue line indicated a threshold for genome-wide alpha of < 0.05 (-log10(p) > 5.58). The effect plots for the top SNP are shown on the right, minor allele frequency (MAF) indicated.

*Reep3* is predicted to play a role in tubular network organization. It is expressed in cochlea (Rousset et al., 2020, Liu et al. 2018, Kolla et al. 2020), but was not detected in 10 month old CFW mice, and is not known to be associated with hearing loss in any species. *Reep3* has a missense variant in CFW mice in high LD with the top SNP of the QTL (c50T>C; Phe17>Ser, r^2^ with the top SNP 0.911).

*Jmjd1c* is *a* predicted histone demethylase and coactivator for transcription factors. It is expressed in inner and outer hair cells in embryonic and young mice (Liu et al. 2018, Kolla et al. 2020, Elcon et al. 2015), but was not detected in 10 month old CFW mice (Boussaty et al. 2023), and is not known to be associated with hearing loss in any species.

<u>QTL on chromosome 4 at 156 Mb</u> (genetic report in **Supplemental Materials**). This chromosomal region contains QTLs for hearing loss at 10 months for 24 kHz. The top SNP for this QTL also shows an association with hearing loss at 10 months for 32 kHz (-log10(p)= 4.89). This region contains 6 genes: *B3galt6* (UDP-Gal:betaGal beta 1,3-galactosyltransferase, polypeptide 6), *Sdf4* (stromal cell derived factor 4), *Tnfrsf4* (TNF Receptor Superfamily Member 4), *Tnfrsf18* (TNF Receptor Superfamily Member 18), Ttll10 (tubulin tyrosine ligase-like family, member 10), and *Gm16008* (predicted long non-coding RNA). None of these genes are expressed in 10 month old CFW mice (Boussaty et al. 2023), although they are expressed at various levels in cochlea of E16, P0, P1, P7, P16 mice (Rousset et al. 2020; Waldhaus et al. 2015; Liu et al. 2018; Kolla et al. 2020; Elcon et al. 2015; Cai et al. 2015). None of these genes were known to be associated with hearing loss in any other species.

We detected three QTLs for elevated ABR thresholds.

<u>QTL on chromosome 5 around 24 Mb</u> (**Figure 7A**). This QTL for elevated ABR threshold at 6 months for 8 kHz encompasses a >1 Mb chromosomal region containing 35 genes. SNPs in this QTL also show association with hearing loss at 6 months for 4 kHz, and at 10 months for 4 kHz and 24 kHz, although these associations do not reach the significance threshold (-log10(p)= 5.05, 4.84, and 4.11, respectively). This QTL contains several genes that have previously been known to affect hearing. *Asic3* (acid-sensing ion channel 3) is expressed in sensory neurons and participates in neuronal mechanotransduction (Lin et al. 2016; Chuang et al. 2022; Cai et al. 2015), and is expressed in supporting cells in adult CBA/J mice (Liu et al. 2018). Knocking out this gene is known to disrupt hearing in mice (Wu et al. 2009). *Asic3* expression was not detected in 10 month old CFW mice (Boussaty et al. 2023).

**Figure 7.**
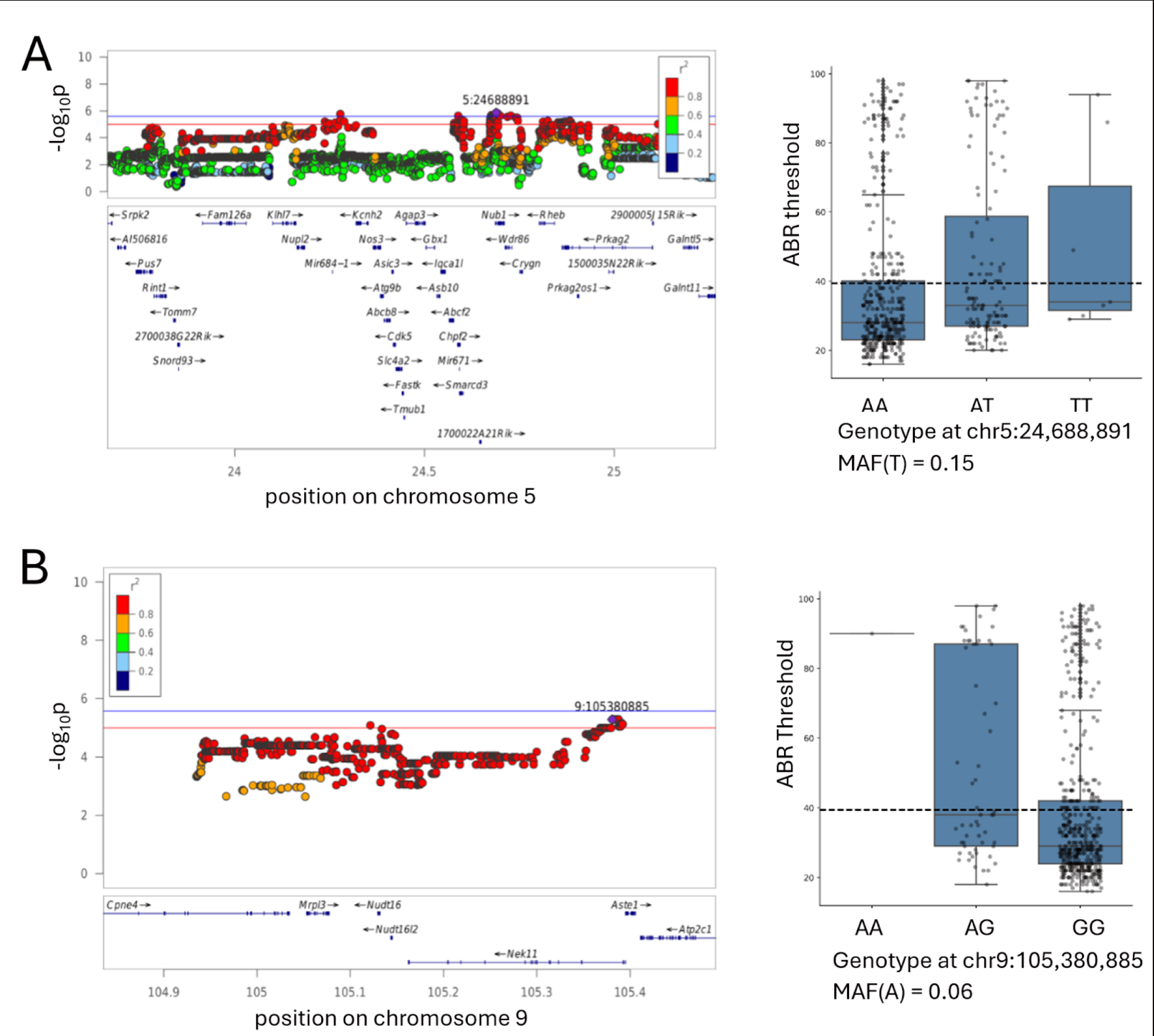
Regional association plot for the ABR threshold at 6 months for 8 kHz.The x-axis shows the position on a chromosome (in Mb); the y-axis shows the significance of the association (-log10 p-value). The individual points represent SNPs. The SNP with the lowest p-value (“top SNP”) is highlighted in purple. The colors represent the correlation between the topSNP and the other SNP. Box plots are showing ABR thresholds in each animal grouped by the genotype at the top SNP, minor allele frequency (MAF) indicated.

*Slc4a2* (solute carrier family 4 (anion exchanger), member 2) plays a role in ion homeostasis, and is expressed in the inner ear (Hosoya et al. 2016; Liu et al. 2018). Knockout mice have multiple severe phenotypes, including deafness (Gawenis et al., 2004). *Slc4a2* expression was not detected in 10 month old CFW mice (Boussaty et al. 2023).

*Crygn* (crystallin, gamma N) is expressed in newborn (Cai et al. 2015) and adult mice (Liu et al. 2018), its expression is required for post migratory survival and proper function of auditory hindbrain neurons; ablation of this gene does not affect ABR thresholds but causes an increase in the amplitude of wave IV (Hartwich et al. 2016). *Crygn* expression was not detected in 10 month old CFW mice (Boussaty et al. 2023).

*Nos3* (nitric oxide synthase 3, endothelial cell) is expressed in inner and outer hair cells of newborn (Cai et al. 2015) and adult mice ((Liu et al. 2018). In humans, polymorphisms in NOS3 are associated with sudden sensorineural hearing loss (Kitoh et al. 2017). *Nos3* expression was not detected in 10 month old CFW mice (Boussaty et al. 2023).

*Cdk5* (cyclin-dependent kinase 5) is expressed ubiquitously in the cochlea of newborn (Elkon et al. 2015; Cai et al. 2015) and adult mice (Zhai et al. 2018; Liu et al. 2018). The cochlea-specific inactivation of Cdk5 causes hearing loss in mice due to loss of stereocilia (Zhai et al. 2018). *Cdk5* expression was not detected in 10 month old CFW mice (Boussaty et al. 2023).

The other genes located in this QTL were not previously reported to be associated with hearing loss. Only one of them, *Prkag2* (protein kinase AMP-activated non-catalytic subunit gamma 2) is expressed in 10 month old CFW mice, mostly in hair cells and spiral ganglion neurons (Boussaty et al. 2023). Due to the number of genes in this interval, it is unclear which one might be the causal gene in this population. However, our work provides support for the role of *Prkag2* in hearing loss (see below).

<u>QTL on chromosome 9 around 105 Mb</u> (**Figure 7B**). This chromosomal region contains QTLs for elevated ABR threshold at 6 months for 8 kHz. It contains 6 genes: *Cpne4* (copine IV), *Mrpl3* (mitochondrial ribosomal protein L3), *Nudt16* (nudix hydrolase 16), *Nek11* (NIMA (never in mitosis gene a)-related expressed kinase 11), and *Aste1* (asteroid homolog 1).

*Cpne4* is expressed in supporting cells in newborn mice (Cai et al. 2015) and at low levels in pillar cells in adult mice (Liu et al. 2018). In 10 month old CFW mice, *Cpne4* expression was detected in spiral ganglion neurons (Boussaty et al. 2023). Copine family of Ca-dependent membrane adaptors are well studied in retinal ganglion cells (Goel et al. 2019; Goel et al. 2021). This study raise a possibility that *Cpna4* is also involved in functioning of spiral ganglion cells in cochlea, and associated with hearing loss.

*Mrpl3* is expressed in cochlear cells of newborn (Elkon et al. 2015; Cai et al. 2015) and adult mice (Liu et al. 2018). The expression is not detected in 10 month old CFW mice (Boussaty et al. 2023). CFW mice have a missense mutation in *Mrpl3* (c55G>A; Ala19Thr, r^2^ with the top SNP 0.822). Mutations in *Mrpl3* have been previously reported to cause altered ribosome assembly and abnormal function of respiratory chain complexes (Bursle et al. 2017).

*Nudt16* is expressed in the cochlea of newborn (Kolla et al. 2020; Elkon et al. 2015; Cai et al. 2015) and adult mice (Liu et al. 2018). It was not detected in 10 month old CFW mice (Boussaty et al. 2023). CFW mice have a missense mutation in *Nudt16* (c457C>A; Val153Met, r^2^ with the top SNP 0.911).

*Nek11* regulates cell cycle. It is expressed in hair cells in newborn and adult mice (Kolla et al. 2020; Elkon et al. 2015; Cai et al. 2015). In 10 month old CFW mice it is detected in hair cells at low level, but mostly expressed in a novel cell type characterized by expression of *Dnah12* and *Rgs22* (Boussaty et al. 2023). *Dnah11*, a candidate gene discussed above, is also expressed in this novel cell type, providing an intriguing possibility of the role of this novel cell type in hearing loss.

*Aste1* is expressed mostly in hair cells in newborn and adult mice (Kolla et al. 2020; Elkon et al. 2015; Cai et al. 2015). It is not detected in 10 month old CFW mice (Boussaty et al. 2023).

<u>QTL on chromosome 11 around 16 Mb</u> (genetic report in **Supplemental Materials**). This QTL is associated with elevated ABR thresholds at 10 months at 32 kHz. It contains two genes: *Vstm2a* (V-set and transmembrane domain containing 2A) and *Sec61g* (SEC61 translocon subunit gamma).

*Vstm2a* and *Sec61g* are expressed in supporting cells of newborn mice (Elkon et al. 2015’ Cai et al. 2015), but were not detected in 10 month old CFW mice (Boussaty et al. 2023). We are not aware of any prior associations between these genes and any hearing related phenotypes.

Prkag2 deficiency causes Age-related hearing loss at high frequency.

We constructed Prkag2 constitutive null mice, and aged these animals, along with the littermate controls, in an effort to assess the impact of Prkag2 deficiency on hearing. The onset of high frequency hearing loss at 20 weeks in the mutants in comparison to the wild-type (**Figure 8**) and the loss of not only outer hair cells, as expected on the C57BL/6 background, but inner hair cells at 2 years (**Figure 8**) confirms a role for Prkag2 in hearing.

**Figure 8.**
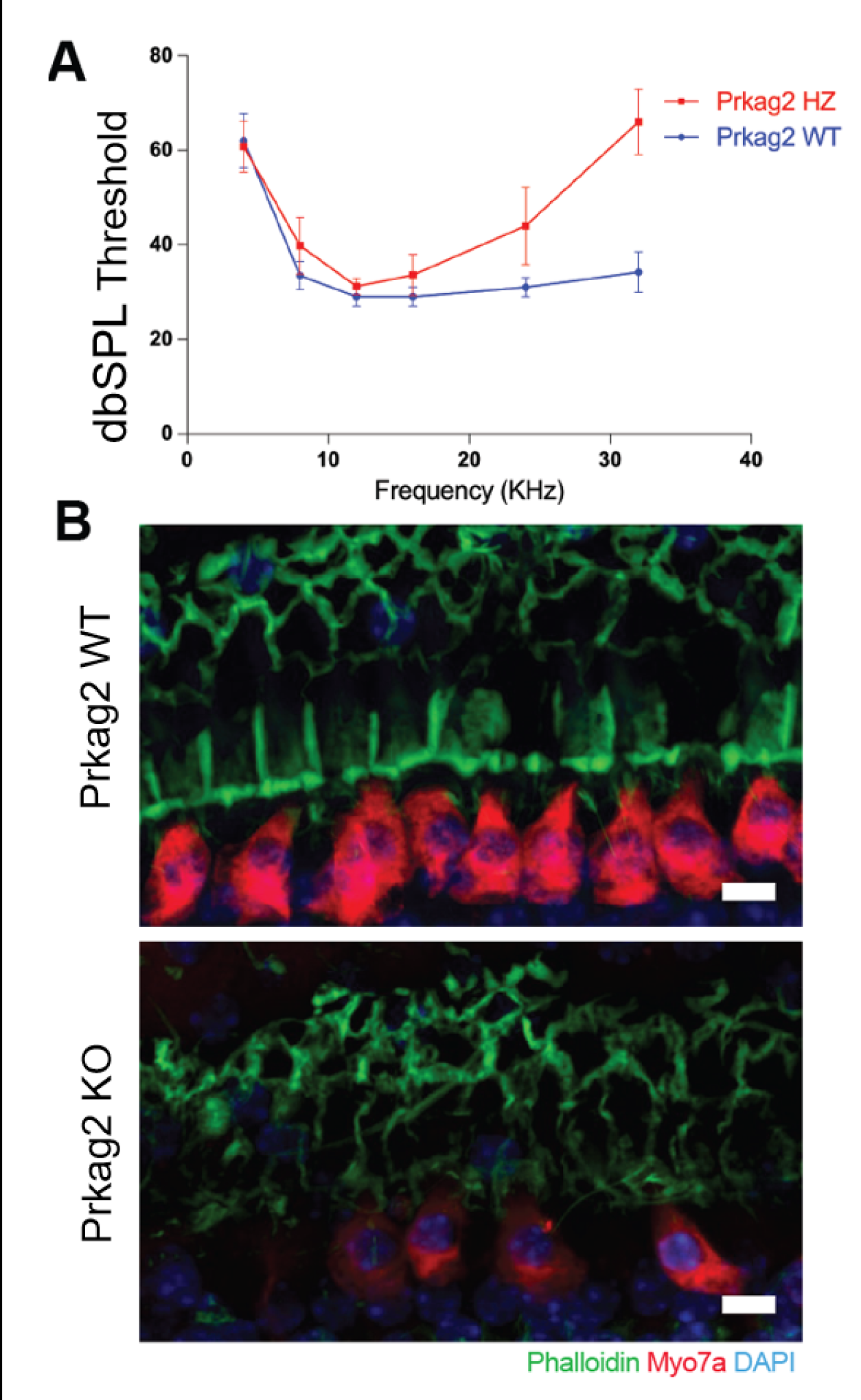
Prkag2-deficient mice show increased sensitivity to ARHL. **A**. Prkag2-deficient mice show high frequency hearing loss as measured by ABR thresholds in 20 month old mice, N = 5. **B**. Confocal images from cochkea samples of 2 years old mice shaw that wild type littermates have higher IHC preservation in comparison to Prkag2-deficient mice. Scale bar is 10 micorometers.

## Discussion

ARHL is a complex trait, meaning that is influenced by many genetic variants, each having a small effect size. In humans, 153 genes that are associated with hearing loss were described as of 2024 (Walls et al. 2024), with various effect sizes and various degrees of confidence. However, this list is not exhaustive and the genetic architecture of hearing loss remains an active field of discovery. Animal models allow gene discovery, which can help to refine our understanding of biological pathways that contribute to hearing loss. In the current work, we use CFW mice to find the genetic underpinnings of age-related hearing loss. The variability in hearing loss in CFW mice has been reported previously, where a subset of mice did not respond to the 120 dB pulses at the age around 4 month old (Parker et al. 2016), in agreement with the current report. This strain was not previously used for genetic studies of hearing loss, therefore we are taking advantage of the genetic variation present in this strain. The outbred nature of the population allows for precise mapping meaning that identified loci often contain a small number of genes.

The SNP heritability of the traits ranged from 0 to 0.42 (**Table 2**). Unlike heritability estimates from inbred strain panels or twins designs, SNP heritability is expected to be lower, since it measures the proportion of phenotypic variance explained by all measured SNPs (Yang et al. 2017).

Heritability of hearing thresholds reported in human studies varies from 0.2 to 0.54 in twin studies (Duan et al. 2019), and 0.13 to 0.7 for SNP-based heritability (Schmitz et al. 2021). Similaraly, the heritability of age-related hearing impairment tends to be higher in twin study designs (0 to 0.65; Bogo et al. 2015, Karlsson et al. 1997) than in association studies (0.03 to 0.22; Fransen et al. 2015; Hoffmann et al. 2016; Huyghe et al. 2008; Wells et al. 2019). In animal models, using a panel of BXD mouse strains, heritability was estimated from 0.21 to 0.70, depending on the exact phenotype (Nagtegaal et al. 2012; Zheng et al. 2020).

We examined the genetic correlations between all possible pairs of traits (**Fig. 3**). Within each age, the correlations between frequencies tend to be higher, forming a characteristic “triangle”. The correlation between ABR thresholds measured at 1 month and ABR threshold or deafness at 6 and 10 months were lower, suggesting that the genetic underpinnings of hearing loss in older mice is less similar to that observed in younger mice.

We discovered 10 loci associated with 7 ARHL traits. Due to the small size of most of the implicated regions (**Fig. 2B**), the QTLs contained from 0 to 35 genes (**Table 3**).

The most interesting genes are those with known coding variation, those that are expressed in cochlea tissue based on several datasets publicly available in gEAR portal (Boussaty et al. 2023; Kolla et al. 2020; Elkon et al. 2015; Cai et al. 2015; Waldhaus et al. 2015; Liu et al. 2018) and those that are supported by previous publications. We confirmed the role of *Prkag2* in hearing loss by demonstrating the oncet of high-frequency hearing loss in 20-month old Prkag2 constitutive null mice, comparing to the wild type littermates accompanied by the loss od both inner hair cells and out hair cells. In the future we hope to examine eQTLs in the cochlea of CFW mice, which would offer an additional line of evidence not available in the current analysis.

Previous mouse studies identified multiple loci related to hearing and hearing loss. In particular, many inbred and outbred mice experience very early hearing loss resulting in severe hearing loss as early as 9 weeks (Zheng et al. 1999). In contrast, few CFW mice showed deafness by 4-6 weeks of age, the first time point, when hearing has been measured. However, by the 6 month time point, 30-43% of mice were categorized as being deaf at different frequencies, demonstrating the presence of alleles that cause relatively early onset of deafness. The current study replicates some of those findings. The lack of replication for other loci may simply reflect the fact that CFW mice do not segregate the same variants as other populations, or could be due to insufficient sample size, or type I and type II errors in the current or prior studies.

Previous studies of genetics of hearing loss in mice were focused mostly on the early onset hearing loss. Many mouse strains are homozygous for an *ahl* locus, corresponding to the Cdh23^753A^ variant (SNP rs257098870) which causes early hearing loss (Johnson et al. 2000; Zheng et al. 1999). The position of this variant corresponds to the last nucleotide of exon 7 in GenBank sequence AF308939 and to the last nucleotide of exon 9 in Ref Seq NM_023370 and NM_001252635. The mechanism is mediated by the effect of this SNP on splicing: G at this position results in normal exon splicing, whereas A disrupts the donor splice site sequence and causes in-frame exon skipping (Noben-Trauth et al. 2003). The Cdh23^753G^ allele is associated with resistance to ARHL and is dominant to the recessive Cdh23^753A^ allele, which is associated with ARHL susceptibility. CFW mice carry the Cdh23^753A^ variant at ∼0.63 allele frequency. This was estimated by using a subset of 86 mice that had at least 3 sequence reads spanning rs257098870, enabling the genotype for rs257098870 to be called by bcftools without imputation. Using these data, we confirmed that homozygosity for Cdh23^753A^ significantly increases susceptibility to age-related hearing loss in our cohort (data not shown). However our imputaton-based genotyping strategy was not able to reliably genotype the region around rs257098870. In CFW this region appears to have structural variation that does not align with the reference genome (data not shown). However, this variant is not the only cause of age-related hearing loss, and our other results remain valid, despite our current inability to accurately genotype rs257098870.

Our genetic analysis in CFW mice did not find several other QTLs that have been previously reported in other mouse strains. The regions corresponding to *ahl3* and *ahl6* have not been narrowed down to a specific gene or a variant, but we were not able to reliably call genotypes for *ahl3* and *ahl6* genomic regions, have a similar issue as ahl locus: we did not call genotypes in this region, most likely because there is no variability in these regions in CFW population, as has been shown previously (Parker et al. 2016). *ahl3* (chromosome 17, 67.2 Mb) was discovered in consomic C57BL/6J and MSM mice (Nemoto et al. 2004); *ahl6* (chromosome 18, 44.2 cM) was discovered in outbred Black Swiss mice (Drayton and Noben-Trauth, 2006). Chromosomal regions corresponding to other previously reported loci were both polymorphic and successfully genotyped, therefore we can confidently report that we did not detect hearing loss in CFW mice that is associated with the following loci: *ahl2* (chromosome 5, 79.6 Mb), discovered in C57BL/6J * NOD/LtJ cross (Johnson,and Zheng, 2002, Ohlemiller et al. 2008), *ahl5* (chromosome 10, 81.1 Mb), corresponding to gene Gipc3 and discovered in outbred Black Swiss mice (Drayton and Noben-Trauth, 2006), *ahl8* (chromosome 11, 120 Mb), corresponding to gene Fscn2 and discovered in BXD mice mice (Johnson et al. 2008), M5Ahl8 (chromosome 5, approximately 78-118 Mb) discovered in BXD mice (Johnson et al. 2015) and *ahp* (chromosome 16) discovered in BXD mice (Zheng et al. 2020). It is not surprising that we do not detect QTLs found in other populations. The main reason is that the two causal alleles might not be polymorphic in CFW population; but it is also possible that the original finding was a false positive or our inability to find a QTL could be a false negative or insufficient sample size.

This study is not without limitations. The number of discoveries in a GWAS is dictated by sample size. In this study, the largest sample sizes were obtained at the 6 month time point. We had originally planned to use a 14 month time point but found that a significant fraction of mice did not live long enough, thus, the 10 month time point was introduced after the study was initiated, and so has fewer subjects. In addition, because a signifncat number of mice were deaf by the 6 and 10 month time points, they could not be used for the analysis of ABR threshold. Another limitation of this study is that we did not explore the onset of deafness, which occurred in 30-43% of mice (varies for different frequencies) sometime between the 1 and 6 month time points. Future studies that examine this process could yield additional insights. Finally, our analysis of genes within implicated regions accounted for coding polymorphisms, but did not examine eQTLs because no eQTL data for the cochlea of CFW mice are available. We plan to develop such data in the future.

In conclusions, we performed a GWAS for ARHL traits using 946 CFW outbred mice - a population previously not used in hearing loss studies. We identified 10 QTLs that offer new insights into genetic underpinning of this pathology, identifying novel candidate genes, including *Dnah11*, *Rapgef5*, *Cpne4*, *Prkag2*, and *Nek11*. Using constitutive knockout mouse model, we confirmed that *Prkag2* plays a role in age-related hearing loss. Other candidate genes identified in this and future studies can be manipulated to explore their role in hearing loss.

Another important future direction will be to explore expression of the candidate genes in spatial manner to better identify cells and structures that are affected.

## Acknowledgments

This publication includes data generated at the UC San Diego IGM Genomics Center utilizing an Illumina NovaSeq 6000 and NovaSeq X Plus that were purchased with funding from a National Institutes of Health SIG grant (#S10 OD026929).

## Supplemental materials

1. The Genetic Analysis Report https://palmerlab.s3.sdsc.edu/tsanches_dash_genotypes/gwas_results/r01_friedman_arhl_2020/results /gwas_report.html
2. **Supplemental Figure 1.** **Supplemental Figure 1.**
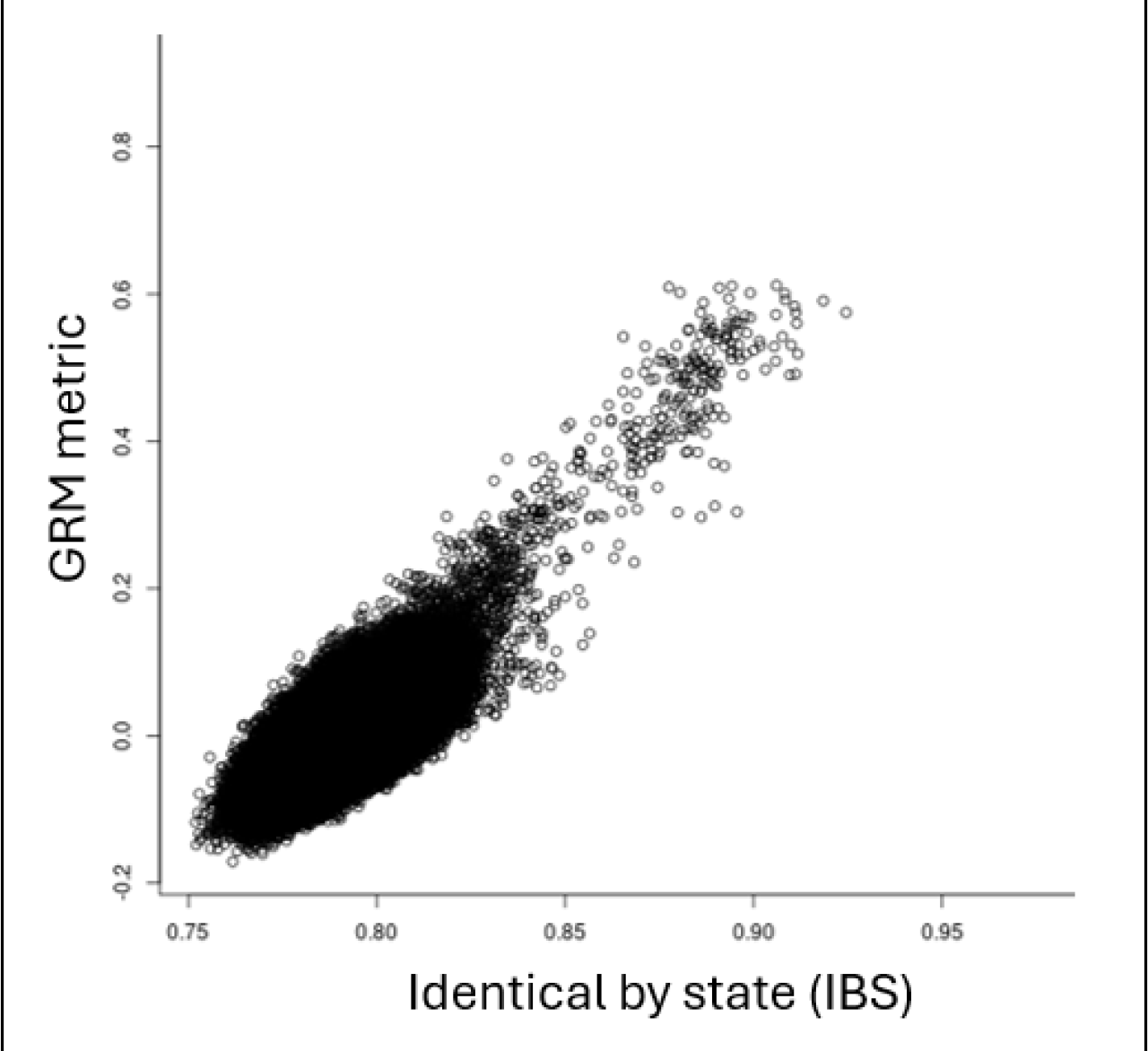
Scatterplot of relatedness expressed as Identical By State (IBS) score and GRM metric shows that there is a group of pairs with high relatedness. There are 252 unique mice in the group that have IBS >0.87 and GRM >0.35, most likely siblings or close cousins.

## References

1. Bainbridge KE, Wallhagen MI. Hearing loss in an aging American population: extent, impact, and management. Annu Rev Public Health. 2014;35:139–52. doi: 10.1146/annurev-publhealth-032013-182510. PMID: 24641557.

2. Belanger JM, Heinonen T, Famula TR, Mandigers PJJ, Leegwater PA, Hytönen MK, Lohi H, Oberbauer AM. Validation of a Chromosome 14 Risk Haplotype for Idiopathic Epilepsy in the Belgian Shepherd Dog Found to Be Associated with an Insertion in the RAPGEF5 Gene. Genes (Basel). 2022 Jun 23;13(7):1124. doi: 10.3390/genes13071124. PMID: 35885906; PMCID: PMC9323784.

3. Blake JA, Baldarelli R, Kadin JA, Richardson JE, Smith CL, Bult CJ; Mouse Genome Database Group. 2021. Mouse Genome Database (MGD): Knowledgebase for mouse-human comparative biology. Nucleic Acids Res. 2021 Jan 8;49(D1):D981-D987

4. Bogo R, Farah A, Johnson AC, Karlsson KK, Pedersen NL, Svartengren M, Skjönsberg Å. The role of genetic factors for hearing deterioration across 20 years: a twin study. J Gerontol A Biol Sci Med Sci. 2015 May;70(5):647–53. doi: 10.1093/gerona/glu245. Epub 2015 Feb 9. PMID: 25665831.

5. Boussaty EC, Ninoyu Y, Andrade L, Li Q, Takeya R, Sumimoto H, Ohyama T, Wahlin KJ, Manor U, Friedman RA. Altered Fhod3 Expression Involved in Progressive High-Frequency Hearing Loss via Dysregulation of Actin Polymerization Stoichiometry in The Cuticular Plate. bioRxiv [Preprint]. 2023 Aug 27:2023.07.20.549974. doi: 10.1101/2023.07.20.549974. Update in: PLoS Genet. 2024 Mar 18;20(3):e1011211. PMID: 37546952; PMCID: PMC10401921.

6. Boussaty EC, Tedeschi N, Novotny M, Ninoyu Y, Du E, Draf C, Zhang Y, Manor U, Scheuermann RH, Friedman R. Cochlear transcriptome analysis of an outbred mouse population (CFW). Front Cell Neurosci. 2023 Nov 29;17:1256619. doi: 10.3389/fncel.2023.1256619. PMID: 38094513; PMCID: PMC10716316.

7. Bok J, Chang W, Wu DK. Patterning and morphogenesis of the vertebrate inner ear. Int J Dev Biol. 2007;51(6-7):521–33. doi: 10.1387/ijdb.072381jb. PMID: 17891714.

8. Bowl MR, Simon MM, Ingham NJ, Greenaway S, Santos L, Cater H, Taylor S, Mason J, Kurbatova N, Pearson S, Bower LR, Clary DA, Meziane H, Reilly P, Minowa O, Kelsey L; International Mouse Phenotyping Consortium; Tocchini-Valentini GP, Gao X, Bradley A, Skarnes WC, Moore M, Beaudet AL, Justice MJ, Seavitt J, Dickinson ME, Wurst W, de Angelis MH, Herault Y, Wakana S, Nutter LMJ, Flenniken AM, McKerlie C, Murray SA, Svenson KL, Braun RE, West DB, Lloyd KCK, Adams DJ, White J, Karp N, Flicek P, Smedley D, Meehan TF, Parkinson HE, Teboul LM, Wells S, Steel KP, Mallon AM, Brown SDM. A large scale hearing loss screen reveals an extensive unexplored genetic landscape for auditory dysfunction. Nat Commun. 2017 Oct 12;8(1):886. doi: 10.1038/s41467-017-00595-4. PMID: 29026089; PMCID: PMC5638796.

9. Browning BL, Browning SR. Genotype Imputation with Millions of Reference Samples. Am J Hum Genet. 2016 Jan 7;98(1):116–26. doi: 10.1016/j.ajhg.2015.11.020. PMID: 26748515; PMCID: PMC4716681.

10. Bursle C, Narendra A, Chuk R, Cardinal J, Justo R, Lewis B, Coman D. COXPD9 an Evolving Multisystem Disease; Congenital Lactic Acidosis, Sensorineural Hearing Loss, Hypertrophic Cardiomyopathy, Cirrhosis and Interstitial Nephritis. JIMD Rep. 2017;34:105–109. doi: 10.1007/8904_2016_13. Epub 2016 Nov 5. PMID: 27815843; PMCID: PMC5509545.

11. Cai T, Jen HI, Kang H, Klisch TJ, Zoghbi HY, Groves AK. Characterization of the transcriptome of nascent hair cells and identification of direct targets of the Atoh1 transcription factor. J Neurosci. 2015 Apr 8;35(14):5870–83. doi: 10.1523/JNEUROSCI.5083-14.2015. PMID: 25855195; PMCID: PMC4388939.

12. Cheng R, Parker CC, Abney M, Palmer AA. Practical considerations regarding the use of genotype and pedigree data to model relatedness in the context of genome-wide association studies. G3 (Bethesda). 2013 Oct 3;3(10):1861-7. doi: 10.1534/g3.113.007948. PMID: 23979941; PMCID: PMC3789811.

13. Cheng R, Palmer AA. A simulation study of permutation, bootstrap, and gene dropping for assessing statistical significance in the case of unequal relatedness. Genetics. 2013 Mar;193(3):1015–8. doi: 10.1534/genetics.112.146332. Epub 2012 Dec 24. PMID: 23267053; PMCID: PMC3583989.

14. Chuang YC, Chen CC. Force From Filaments: The Role of the Cytoskeleton and Extracellular Matrix in the Gating of Mechanosensitive Channels. Front Cell Dev Biol. 2022 May 2;10:886048. doi: 10.3389/fcell.2022.886048. PMID: 35586339; PMCID: PMC9108448.

15. Cingolani P, Platts A, Wang le L, Coon M, Nguyen T, Wang L, Land SJ, Lu X, Ruden DM. A program for annotating and predicting the effects of single nucleotide polymorphisms, SnpEff: SNPs in the genome of Drosophila melanogaster strain w1118; iso-2; iso-3. Fly (Austin). 2012 Apr-Jun;6(2):80-92. doi: 10.4161/fly.19695. PMID: 22728672; PMCID: PMC3679285.

16. Davies, Robert W., Jonathan Flint, Simon Myers, and Richard Mott. 2016. “Rapid Genotype Imputation from Sequence without Reference Panels.” Nature Genetics 48 (8): 965–69.

17. De Angelis F, Zeleznik OA, Wendt FR, Pathak GA, Tylee DS, De Lillo A, Koller D, Cabrera-Mendoza B, Clifford RE, Maihofer AX, Nievergelt CM, Curhan GC, Curhan SG, Polimanti R. Sex differences in the polygenic architecture of hearing problems in adults. Genome Med. 2023 May 11;15(1):36. doi: 10.1186/s13073-023-01186-3. PMID: 37165447; PMCID: PMC10173489.

18. Deal JA, Betz J, Yaffe K, Harris T, Purchase-Helzner E, Satterfield S, Pratt S, Govil N, Simonsick EM, Lin FR; Health ABC Study Group. Hearing Impairment and Incident Dementia and Cognitive Decline in Older Adults: The Health ABC Study. J Gerontol A Biol Sci Med Sci. 2017 May 1;72(5):703–709. doi: 10.1093/gerona/glw069. PMID: 27071780; PMCID: PMC5964742.

19. Deal JA, Reed NS, Kravetz AD, Weinreich H, Yeh C, Lin FR, Altan A. Incident Hearing Loss and Comorbidity: A Longitudinal Administrative Claims Study. JAMA Otolaryngol Head Neck Surg. 2019 Jan 1;145(1):36–43. doi: 10.1001/jamaoto.2018.2876. PMID: 30419134; PMCID: PMC6439817.

20. Drayton M, Noben-Trauth K. Mapping quantitative trait loci for hearing loss in Black Swiss mice. Hear Res. 2006 Feb;212(1-2):128–39. doi: 10.1016/j.heares.2005.11.006. Epub 2006 Jan 19. PMID: 16426780.

21. Du EY, Boussaty EC, La Monte OA, Dixon PR, Zhou TY, Friedman RA. Large-scale phenotyping and characterization of age-related hearing loss in outbred CFW mice. Hear Res. 2022 Oct;424:108605. doi: 10.1016/j.heares.2022.108605. Epub 2022 Sep 5. PMID: 36088865.

22. Duan H, Zhang D, Liang Y, Xu C, Wu Y, Tian X, Pang Z, Tan Q, Li S, Qiu C. Heritability of Age-Related Hearing Loss in Middle-Aged and Elderly Chinese: A Population-Based Twin Study. Ear Hear. 2019 Mar/Apr;40(2):253-259. doi: 10.1097/AUD.0000000000000610. PMID: 29794565.

23. Elkon R, Milon B, Morrison L, Shah M, Vijayakumar S, Racherla M, Leitch CC, Silipino L, Hadi S, Weiss-Gayet M, Barras E, Schmid CD, Ait-Lounis A, Barnes A, Song Y, Eisenman DJ, Eliyahu E, Frolenkov GI, Strome SE, Durand B, Zaghloul NA, Jones SM, Reith W, Hertzano R. RFX transcription factors are essential for hearing in mice. Nat Commun. 2015 Oct 15;6:8549. doi: 10.1038/ncomms9549. PMID: 26469318; PMCID: PMC4634137.

24. Fu X, An Y, Wang H, Li P, Lin J, Yuan J, Yue R, Jin Y, Gao J, Chai R. Deficiency of Klc2 Induces Low-Frequency Sensorineural Hearing Loss in C57BL/6 J Mice and Human. Mol Neurobiol. 2021 Sep;58(9):4376–4391. doi: 10.1007/s12035-021-02422-w. Epub 2021 May 20. PMID: 34014435.

25. Gawenis LR, Ledoussal C, Judd LM, Prasad V, Alper SL, Stuart-Tilley A, Woo AL, Grisham C, Sanford LP, Doetschman T, Miller ML, Shull GE. Mice with a targeted disruption of the AE2 Cl-/HCO3- exchanger are achlorhydric. J Biol Chem. 2004 Jul 16;279(29):30531–9. doi: 10.1074/jbc.M403779200. Epub 2004 Apr 30. PMID: 15123620.

26. Goel M, Aponte AM, Wistow G, Badea TC. Molecular studies into cell biological role of Copine-4 in Retinal Ganglion Cells. PLoS One. 2021 Nov 30;16(11):e0255860. doi: 10.1371/journal.pone.0255860. PMID: 34847148; PMCID: PMC8631636.

27. Goel M, Li T, Badea TC. Differential expression and subcellular localization of Copines in mouse retina. J Comp Neurol. 2019 Oct 1;527(14):2245–2262. doi: 10.1002/cne.24684. Epub 2019 Mar 28. PMID: 30866042; PMCID: PMC6656606.

28. Johnson KR, Longo-Guess CM, Gagnon LH. A QTL on Chr 5 modifies hearing loss associated with the fascin-2 variant of DBA/2J mice. Mamm Genome. 2015 Aug;26(7-8):338–47. doi: 10.1007/s00335-015-9574-y. Epub 2015 Jun 20. PMID: 26092689; PMCID: PMC4629822.

29. Johnson KR, Longo-Guess CM, Gagnon LH. A QTL on Chr 5 modifies hearing loss associated with the fascin-2 variant of DBA/2J mice. Mamm Genome. 2015 Aug;26(7-8):338–47. doi: 10.1007/s00335-015-9574-y. Epub 2015 Jun 20. PMID: 26092689; PMCID: PMC4629822.

30. Johnson KR, Longo-Guess C, Gagnon LH, Yu H, Zheng QY. A locus on distal chromosome 11 (ahl8) and its interaction with Cdh23 ahl underlie the early onset, age-related hearing loss of DBA/2J mice. Genomics. 2008 Oct;92(4):219–25. doi: 10.1016/j.ygeno.2008.06.007. Epub 2008 Aug 15. PMID: 18662770; PMCID: PMC2836023.

31. Johnson KR, Zheng QY. Ahl2, a second locus affecting age-related hearing loss in mice. Genomics. 2002 Nov;80(5):461–4. PMID: 12408962; PMCID: PMC2862211.

32. Johnson KR, Zheng QY, Erway LC. A major gene affecting age-related hearing loss is common to at least ten inbred strains of mice. Genomics. 2000 Dec 1;70(2):171–80. doi: 10.1006/geno.2000.6377. PMID: 11112345.

33. Hartwich H, Rosengauer E, Rüttiger L, Wilms V, Waterholter SK, Nothwang HG. Functional Role of γ-Crystallin N in the Auditory Hindbrain. PLoS One. 2016 Aug 12;11(8):e0161140. doi: 10.1371/journal.pone.0161140. PMID: 27517863; PMCID: PMC4982622.

34. Hoffmann TJ, Keats BJ, Yoshikawa N, Schaefer C, Risch N, Lustig LR. A Large Genome-Wide Association Study of Age-Related Hearing Impairment Using Electronic Health Records. PLoS Genet. 2016 Oct 20;12(10):e1006371. doi: 10.1371/journal.pgen.1006371. PMID: 27764096; PMCID: PMC5072625.

35. Hosoya M, Fujioka M, Kobayashi R, Okano H, Ogawa K. Overlapping expression of anion exchangers in the cochlea of a non-human primate suggests functional compensation. Neurosci Res. 2016 Sep;110:1–10. doi: 10.1016/j.neures.2016.04.002. Epub 2016 Apr 19. PMID: 27091614.

36. Huyghe JR, Van Laer L, Hendrickx JJ, Fransen E, Demeester K, Topsakal V, Kunst S, Manninen M, Jensen M, Bonaconsa A, Mazzoli M, Baur M, Hannula S, Mäki-Torkko E, Espeso A, Van Eyken E, Flaquer A, Becker C, Stephens D, Sorri M, Orzan E, Bille M, Parving A, Pyykkö I, Cremers CW, Kremer H, Van de Heyning PH, Wienker TF, Nürnberg P, Pfister M, Van Camp G. Genome-wide SNP-based linkage scan identifies a locus on 8q24 for an age-related hearing impairment trait. Am J Hum Genet. 2008 Sep;83(3):401–7. doi: 10.1016/j.ajhg.2008.08.002. Epub 2008 Aug 28. PMID: 18760390; PMCID: PMC2556434.

37. Fransen E, Bonneux S, Corneveaux JJ, Schrauwen I, Di Berardino F, White CH, Ohmen JD, Van de Heyning P, Ambrosetti U, Huentelman MJ, Van Camp G, Friedman RA. Genome-wide association analysis demonstrates the highly polygenic character of age-related hearing impairment. Eur J Hum Genet. 2015 Jan;23(1):110–5. doi: 10.1038/ejhg.2014.56. Epub 2014 Jun 18. PMID: 24939585; PMCID: PMC4266741.

38. Friedman RA, Van Laer L, Huentelman MJ, Sheth SS, Van Eyken E, Corneveaux JJ, Tembe WD, Halperin RF, Thorburn AQ, Thys S, Bonneux S, Fransen E, Huyghe J, Pyykkö I, Cremers CW, Kremer H, Dhooge I, Stephens D, Orzan E, Pfister M, Bille M, Parving A, Sorri M, Van de Heyning PH, Makmura L, Ohmen JD, Linthicum FH Jr, Fayad JN, Pearson JV, Craig DW, Stephan DA, Van Camp G. GRM7 variants confer susceptibility to age-related hearing impairment. Hum Mol Genet. 2009 Feb 15;18(4):785–96. doi: 10.1093/hmg/ddn402. Epub 2008 Dec 1. PMID: 19047183; PMCID: PMC2638831.

39. Fu X, An Y, Wang H, Li P, Lin J, Yuan J, Yue R, Jin Y, Gao J, Chai R. Deficiency of Klc2 Induces Low-Frequency Sensorineural Hearing Loss in C57BL/6 J Mice and Human. Mol Neurobiol. 2021 Sep;58(9):4376–4391. doi: 10.1007/s12035-021-02422-w. Epub 2021 May 20. PMID: 34014435.

40. Gileta AF, Fitzpatrick CJ, Chitre AS, St Pierre CL, Joyce EV, Maguire RJ, McLeod AM, Gonzales NM, Williams AE, Morrow JD, Robinson TE, Flagel SB, Palmer AA. Genetic characterization of outbred Sprague Dawley rats and utility for genome-wide association studies. PLoS Genet. 2022 May 31;18(5):e1010234. doi: 10.1371/journal.pgen.1010234. PMID: 35639796; PMCID: PMC9187121.

41. Girotto G, Pirastu N, Sorice R, Biino G, Campbell H, d’Adamo AP, Hastie ND, Nutile T, Polasek O, Portas L, Rudan I, Ulivi S, Zemunik T, Wright AF, Ciullo M, Hayward C, Pirastu M, Gasparini P. Hearing function and thresholds: a genome-wide association study in European isolated populations identifies new loci and pathways. J Med Genet. 2011 Jun;48(6):369–74. doi: 10.1136/jmg.2010.088310. Epub 2011 Apr 14. PMID: 21493956.

42. Gonzales NM, Seo J, Hernandez Cordero AI, St Pierre CL, Gregory JS, Distler MG, Abney M, Canzar S, Lionikas A, Palmer AA. Genome-wide association analysis in a mouse advanced intercross line. Nat Commun. 2018 Dec 4;9(1):5162. doi: 10.1038/s41467-018-07642-8. PMID: 30514929; PMCID: PMC6279738.

43. Kalra G, Milon B, Casella AM, Herb BR, Humphries E, Song Y, Rose KP, Hertzano R, Ament SA. Biological insights from multi-omic analysis of 31 genomic risk loci for adult hearing difficulty. PLoS Genet. 2020 Sep 28;16(9):e1009025. doi: 10.1371/journal.pgen.1009025. PMID: 32986727; PMCID: PMC7544108.

44. Karlsson KK, Harris JR, Svartengren M. Description and primary results from an audiometric study of male twins. Ear Hear. 1997 Apr;18(2):114–20. doi: 10.1097/00003446-199704000-00003. PMID: 9099560.

45. Kitoh R, Nishio SY, Usami SI. Prognostic impact of gene polymorphisms in patients with idiopathic sudden sensorineural hearing loss. Acta Otolaryngol. 2017;137(sup565):S24–S29. doi: 10.1080/00016489.2017.1296971. Epub 2017 Apr 1. PMID: 28366034.

46. Lee SH, Yang J, Goddard ME, Visscher PM, Wray NR. Estimation of pleiotropy between complex diseases using single-nucleotide polymorphism-derived genomic relationships and restricted maximum likelihood. Bioinformatics. 2012 Oct 1;28(19):2540–2. doi: 10.1093/bioinformatics/bts474. Epub 2012 Jul 26. PMID: 22843982; PMCID: PMC3463125.

47. Lewis MA, Nolan LS, Cadge BA, Matthews LJ, Schulte BA, Dubno JR, Steel KP, Dawson SJ. Whole exome sequencing in adult-onset hearing loss reveals a high load of predicted pathogenic variants in known deafness-associated genes and identifies new candidate genes. BMC Med Genomics. 2018 Sep 4;11(1):77. doi: 10.1186/s12920-018-0395-1. PMID: 30180840; PMCID: PMC6123954.

48. Lewis MA, Di Domenico F, Ingham NJ, Prosser HM, Steel KP. Hearing impairment due to Mir183/96/182 mutations suggests both loss and gain of function effects. Dis Model Mech. 2020 Dec 14;14(2):dmm047225. doi: 10.1242/dmm.047225. Epub ahead of print. PMID: 33318051; PMCID: PMC7903918.

49. Lin FR, Albert M. Hearing loss and dementia - who is listening? Aging Ment Health. 2014;18(6):671–3. doi: 10.1080/13607863.2014.915924. PMID: 24875093; PMCID: PMC4075051.

50. Lin SH, Cheng YR, Banks RW, Min MY, Bewick GS, Chen CC. Evidence for the involvement of ASIC3 in sensory mechanotransduction in proprioceptors. Nat Commun. 2016 May 10;7:11460. doi: 10.1038/ncomms11460. PMID: 27161260; PMCID: PMC4866049.

51. Listgarten J, Lippert C, Kadie CM, Davidson RI, Eskin E, Heckerman D. Improved linear mixed models for genome-wide association studies. Nat Methods. 2012 May 30;9(6):525–6. doi: 10.1038/nmeth.2037. PMID: 22669648; PMCID: PMC3597090.

52. Liu H, Chen L, Giffen KP, Stringham ST, Li Y, Judge PD, Beisel KW, He DZZ. Cell-Specific Transcriptome Analysis Shows That Adult Pillar and Deiters’ Cells Express Genes Encoding Machinery for Specializations of Cochlear Hair Cells. Front Mol Neurosci. 2018 Oct 1;11:356. doi: 10.3389/fnmol.2018.00356. PMID: 30327589; PMCID: PMC6174830.

53. Lynch CJ. The so-called Swiss mouse. Lab Anim Care. 1969 Apr;19(2):214–20. PMID: 4240230.

54. Kolla L, Kelly MC, Mann ZF, Anaya-Rocha A, Ellis K, Lemons A, Palermo AT, So KS, Mays JC, Orvis J, Burns JC, Hertzano R, Driver EC, Kelley MW. Characterization of the development of the mouse cochlear epithelium at the single cell level. Nat Commun. 2020 May 13;11(1):2389. doi: 10.1038/s41467-020-16113-y. PMID: 32404924; PMCID: PMC7221106.

55. Momi SK, Wolber LE, Fabiane SM, MacGregor AJ, Williams FM. Genetic and Environmental Factors in Age-Related Hearing Impairment. Twin Res Hum Genet. 2015 Aug;18(4):383–92. doi: 10.1017/thg.2015.35. Epub 2015 Jun 17. PMID: 26081266.

56. MGI, E-MTAB-5800, RNA-seq of organ of Corti RNA from mice carrying a null allele of Mir183 and Mir96

57. Nagtegaal AP, Spijker S, Crins TT; Neuro-Bsik Mouse Phenomics Consortium; Borst JG. A novel QTL underlying early-onset, low-frequency hearing loss in BXD recombinant inbred strains. Genes Brain Behav. 2012 Nov;11(8):911–20. doi: 10.1111/j.1601-183X.2012.00845.x. Epub 2012 Oct 8. PMID: 22989164.

58. Nemoto M, Morita Y, Mishima Y, Takahashi S, Nomura T, Ushiki T, Shiroishi T, Kikkawa Y, Yonekawa H, Kominami R. Ahl3, a third locus on mouse chromosome 17 affecting age-related hearing loss. Biochem Biophys Res Commun. 2004 Nov 26;324(4):1283–8. doi: 10.1016/j.bbrc.2004.09.186. PMID: 15504353.

59. Newman DL, Fisher LM, Ohmen J, Parody R, Fong CT, Frisina ST, Mapes F, Eddins DA, Robert Frisina D, Frisina RD, Friedman RA. GRM7 variants associated with age-related hearing loss based on auditory perception. Hear Res. 2012 Dec;294(1-2):125–32. doi: 10.1016/j.heares.2012.08.016. Epub 2012 Oct 25. PMID: 23102807; PMCID: PMC3705704.

60. Nicod, Jérôme, Robert W. Davies, Na Cai, Carl Hassett, Leo Goodstadt, Cormac Cosgrove, Benjamin K. Yee, et al. 2016. “Genome-Wide Association of Multiple Complex Traits in Outbred Mice by Ultra-Low-Coverage Sequencing.” Nature Genetics 48 (8): 912–18.

61. Noben-Trauth K, Zheng QY, Johnson KR. Association of cadherin 23 with polygenic inheritance and genetic modification of sensorineural hearing loss. Nat Genet. 2003 Sep;35(1):21–3. doi: 10.1038/ng1226. Epub 2003 Aug 10. PMID: 12910270; PMCID: PMC2864026.

62. Ohlemiller KK, Rice ME, Gagnon PM. Strial microvascular pathology and age-associated endocochlear potential decline in NOD congenic mice. Hear Res. 2008 Oct;244(1-2):85–97. doi: 10.1016/j.heares.2008.08.001. Epub 2008 Aug 12. PMID: 18727954; PMCID: PMC2630541.

63. Palmer, AA. Abraham A. Palmer Lab Research Data Collection. UC San Diego Library Digital Collections, 2023, 10.6075/j0h13263.

64. Parker, Clarissa C., Shyam Gopalakrishnan, Peter Carbonetto, Natalia M. Gonzales, Emily Leung, Yeonhee J. Park, Emmanuel Aryee, et al. 2016. “Genome-Wide Association Study of Behavioral, Physiological and Gene Expression Traits in Outbred CFW Mice.” Nature Genetics 48 (8): 919–26.

65. Praveen K, Dobbyn L, Gurski L, Ayer AH, Staples J, Mishra S, Bai Y, Kaufman A, Moscati A, Benner C, Chen E, Chen S, Popov A, Smith J; GHS-REGN DiscovEHR collaboration; Regeneron Genetics Center; Decibel-REGN collaboration; Melander O, Jones MB, Marchini J, Balasubramanian S, Zambrowicz B, Drummond MC, Baras A, Abecasis GR, Ferreira MA, Stahl EA, Coppola G. Population-scale analysis of common and rare genetic variation associated with hearing loss in adults. Commun Biol. 2022 Jun 3;5(1):540. doi: 10.1038/s42003-022-03408-7. PMID: 35661827; PMCID: PMC9166757.

66. Pruim RJ, Welch RP, Sanna S, Teslovich TM, Chines PS, Gliedt TP, Boehnke M, Abecasis GR, Willer CJ. LocusZoom: regional visualization of genome-wide association scan results. Bioinformatics. 2010 Sep 15;26(18):2336–7. doi: 10.1093/bioinformatics/btq419. Epub 2010 Jul 15. PMID: 20634204; PMCID: PMC2935401.

67. Rousset F, Nacher-Soler G, Coelho M, Ilmjarv S, Kokje VBC, Marteyn A, Cambet Y, Perny M, Roccio M, Jaquet V, Senn P, Krause KH. Redox activation of excitatory pathways in auditory neurons as mechanism of age-related hearing loss. Redox Biol. 2020 Feb;30:101434. doi: 10.1016/j.redox.2020.101434. Epub 2020 Jan 20. PMID: 32000019; PMCID: PMC7016250.

68. Schmitz J, Abbondanza F, Paracchini S. Genome-wide association study and polygenic risk score analysis for hearing measures in children. Am J Med Genet B Neuropsychiatr Genet. 2021 Jul;186(5):318–328. doi: 10.1002/ajmg.b.32873. Epub 2021 Sep 3. PMID: 34476894.

69. Schwander M, Xiong W, Tokita J, Lelli A, Elledge HM, Kazmierczak P, Sczaniecka A, Kolatkar A, Wiltshire T, Kuhn P, Holt JR, Kachar B, Tarantino L, Müller U. A mouse model for nonsyndromic deafness (DFNB12) links hearing loss to defects in tip links of mechanosensory hair cells. Proc Natl Acad Sci U S A. 2009 Mar 31;106(13):5252–7. doi: 10.1073/pnas.0900691106. Epub 2009 Mar 6. PMID: 19270079; PMCID: PMC2664065.

70. Su Z, Xiong H, Pang J, Lin H, Lai L, Zhang H, Zhang W, Zheng Y. LncRNA AW112010 Promotes Mitochondrial Biogenesis and Hair Cell Survival: Implications for Age-Related Hearing Loss. Oxid Med Cell Longev. 2019 Oct 27;2019:6150148. doi: 10.1155/2019/6150148. PMID: 31781342; PMCID: PMC6855056.

71. Sun G, Zheng Y, Fu X, Zhang W, Ren J, Ma S, Sun S, He X, Wang Q, Ji Z, Cheng F, Yan K, Liu Z, Belmonte JCI, Qu J, Wang S, Chai R, Liu GH. Single-cell transcriptomic atlas of mouse cochlear aging. Protein Cell. 2023 Apr 13;14(3):180–201. doi: 10.1093/procel/pwac058. PMID: 36933008; PMCID: PMC10098046.

72. Van Laer L, Huyghe JR, Hannula S, Van Eyken E, Stephan DA, Mäki-Torkko E, Aikio P, Fransen E, Lysholm-Bernacchi A, Sorri M, Huentelman MJ, Van Camp G. A genome-wide association study for age-related hearing impairment in the Saami. Eur J Hum Genet. 2010 Jun;18(6):685–93. doi: 10.1038/ejhg.2009.234. Epub 2010 Jan 13. PMID: 20068591; PMCID: PMC2987344.

73. Visel A, Thaller C, Eichele G. GenePaint.org: an atlas of gene expression patterns in the mouse embryo. Nucleic Acids Res. 2004 Jan 1;32(Database issue):D552-6. doi: 10.1093/nar/gkh029. PMID: 14681479; PMCID: PMC308763.

74. Vuckovic D, Dawson S, Scheffer DI, Rantanen T, Morgan A, Di Stazio M, Vozzi D, Nutile T, Concas MP, Biino G, Nolan L, Bahl A, Loukola A, Viljanen A, Davis A, Ciullo M, Corey DP, Pirastu M, Gasparini P, Girotto G. Genome-wide association analysis on normal hearing function identifies PCDH20 and SLC28A3 as candidates for hearing function and loss. Hum Mol Genet. 2015 Oct 1;24(19):5655–64. doi: 10.1093/hmg/ddv279. Epub 2015 Jul 17. PMID: 26188009; PMCID: PMC4572074.

75. Waldhaus J, Durruthy-Durruthy R, Heller S. Quantitative High-Resolution Cellular Map of the Organ of Corti. Cell Rep. 2015 Jun 9;11(9):1385–99. doi: 10.1016/j.celrep.2015.04.062. Epub 2015 May 28. PMID: 26027927; PMCID: PMC4465070.

76. Walls WD, Azaiez H, Smith RJH. Hereditary Hearing Loss Homepage. https://hereditaryhearingloss.org, accessed on April 19, 2024

77. Wells HRR, Freidin MB, Zainul Abidin FN, Payton A, Dawes P, Munro KJ, Morton CC, Moore DR, Dawson SJ, Williams FMK. GWAS Identifies 44 Independent Associated Genomic Loci for Self-Reported Adult Hearing Difficulty in UK Biobank. Am J Hum Genet. 2019 Oct 3;105(4):788–802. doi: 10.1016/j.ajhg.2019.09.008. Epub 2019 Sep 26. PMID: 31564434; PMCID: PMC6817556.

78. Wu WL, Wang CH, Huang EY, Chen CC. Asic3(-/-) female mice with hearing deficit affects social development of pups. PLoS One. 2009 Aug 4;4(8):e6508. doi: 10.1371/journal.pone.0006508. PMID: 19652708; PMCID: PMC2714966.

79. Yalcin B, Nicod J, Bhomra A, Davidson S, Cleak J, Farinelli L, Østerås M, Whitley A, Yuan W, Gan X, Goodson M. Commercially available outbred mice for genome-wide association studies. PLoS genetics. 2010 Sep 2;6(9):e1001085.

80. Yang J, Benyamin B, McEvoy BP, Gordon S, Henders AK, Nyholt DR, Madden PA, Heath AC, Martin NG, Montgomery GW, Goddard ME, Visscher PM. Common SNPs explain a large proportion of the heritability for human height. Nat Genet. 2010 Jul;42(7):565–9. doi: 10.1038/ng.608. Epub 2010 Jun 20. PMID: 20562875; PMCID: PMC3232052.

81. Yang J, Lee SH, Goddard ME, Visscher PM. GCTA: a tool for genome-wide complex trait analysis. Am J Hum Genet. 2011 Jan 7;88(1):76–82. doi: 10.1016/j.ajhg.2010.11.011. Epub 2010 Dec 17. PMID: 21167468; PMCID: PMC3014363.

82. Yang J, Zeng J, Goddard ME, Wray NR, Visscher PM. Concepts, estimation and interpretation of SNP-based heritability. Nat Genet. 2017 Aug 30;49(9):1304–1310. doi: 10.1038/ng.3941. PMID: 28854176.

83. Yang J, Zaitlen NA, Goddard ME, Visscher PM, Price AL. Advantages and pitfalls in the application of mixed-model association methods. Nat Genet. 2014 Feb;46(2):100–6. doi: 10.1038/ng.2876. PMID: 24473328; PMCID: PMC3989144.

84. Yu KS, Frumm SM, Park JS, Lee K, Wong DM, Byrnes L, Knox SM, Sneddon JB, Tward AD. Development of the Mouse and Human Cochlea at Single Cell Resolution. bioRxiv 2019. doi: 10.1101/739680

85. Zhai X, Liu C, Zhao B, Wang Y, Xu Z. Inactivation of Cyclin-Dependent Kinase 5 in Hair Cells Causes Hearing Loss in Mice. Front Mol Neurosci. 2018 Dec 11;11:461. doi: 10.3389/fnmol.2018.00461. PMID: 30618612; PMCID: PMC6297389.

86. Zhang Y, Zhang S, Zhou H, Ma X, Wu L, Tian M, Li S, Qian X, Gao X, Chai R. Dync1li1 is required for the survival of mammalian cochlear hair cells by regulating the transportation of autophagosomes. PLoS Genet. 2022 Jun 21;18(6):e1010232. doi: 10.1371/journal.pgen.1010232. PMID: 35727824; PMCID: PMC9249241.

87. Zheng QY, Johnson KR, Erway LC. Assessment of hearing in 80 inbred strains of mice by ABR threshold analyses. Hear Res. 1999 Apr;130(1-2):94–107. doi: 10.1016/s0378-5955(99)00003-9. PMID: 10320101; PMCID: PMC2855304.

88. Zheng QY, Kui L, Xu F, Zheng T, Li B, McCarty M, Sun Z, Zhang A, Liu L, Starlard-Davenport A, Stepanyan R, Hu BH, Lu L. An Age-Related Hearing Protection Locus on Chromosome 16 of BXD Strain Mice. Neural Plast. 2020 Jun 8;2020:8889264. doi: 10.1155/2020/8889264. PMID: 32587610; PMCID: PMC7298343.

89. Zou, Jennifer, Shyam Gopalakrishnan, Clarissa C. Parker, Jerome Nicod, Richard Mott, Na Cai, Arimantas Lionikas, Robert W. Davies, Abraham A. Palmer, and Jonathan Flint. 2022. “Analysis of Independent Cohorts of Outbred CFW Mice Reveals Novel Loci for Behavioral and Physiological Traits and Identifies Factors Determining Reproducibility.” G3 12 (1).

